# Immunosuppressive myeloid cells induce mesenchymal-like breast cancer stem cells by a mechanism involving membrane-bound TGF-β1

**DOI:** 10.1101/2025.02.21.639264

**Authors:** Thomas Boyer, Céline Blaye, Domitille Chalopin, Mathilde Madéry, Jonathan Boucher, Alexandra Moisand, Julie Giraud, Audrey Theodoly-Lannes, Florent Peyraud, Lornella Seeneevassen, Clément Klein, Samy Mebroukine, Sophie Auriol, Assia Chaibi, Atika Zouine, Darya Alizadeh, Gaetan MacGrogan, Baptiste Lamarthée, Bernard Bonnotte, Eric Bonneil, Philippe P. Roux, Christine Varon, Charlotte Domblides, Nicolas Larmonier

**Affiliations:** University of Bordeaux, CNRS, ImmunoConcEpT, UMR5164, 33000 Bordeaux, France; Bergonié Institute, Bordeaux, France; University of Bordeaux, CNRS, IBGC, UMR5095, 33000 Bordeaux, France; Institut de Recherche en Immunologie et en Cancérologie de l’Université de Montréal, Université de Montréal, Québec, Canada; University of Bordeaux, INSERM, BoRdeaux Institute of onCology (BRIC), UMR1312, 33000 Bordeaux, France; ImmuSmol, 33000 Bordeaux, France; University of Bordeaux, INSERM, Flow Cytometry Facility, UAR 3427, INSERM US05, 33000 Bordeaux, France; City of Hope, T Cell Therapeutics Research Labs, Cellular Immunotherapy Center, Department of Hematology and Hematopoietic Cell Transplantation, Duarte, California; University Marie et Louis Pasteur, EFS, INSERM, UMR RIGHT, F-25000 Besançon, France; University of Bourgogne, INSERM, UMR1098, CHU de Dijon, France; Département de pathologie et biologie cellulaire, Faculté de médecine, Université de Montréal, Québec, Canada; Department of Medical Oncology, Hôpital Saint-André, CHU Bordeaux, France

**Author notes:** **Corresponding Authors:** Nicolas Larmonier, Charlotte Domblides, CNRS UMR 5164 ImmunoConcEpT / UF Biology, Biological and Medical Sciences Department, University of Bordeaux, 2 Rue Dr. Hoffman Martinot, Bordeaux, France. CB and TB equally contributed to the study. NL, CD and CV share senior authorship.

**Keywords:** Tumor-promoting Myeloid cells, Immunosuppression, Tumor immune escape, Cancer stemness, mesenchymal cancer stem cells, membrane TGF-β1

## Abstract

Suppressive myeloid cells play a central role in cancer escape from anti-tumor immunity. Beyond their immunosuppressive function, these cells are capable of exerting multiple other pro-tumoral activities, including the promotion of cancer cell survival, invasion and metastasis. The ability of some myeloid subsets to induce cancer stemness has recently emerged. Here we demonstrated that human immunosuppressive myeloid cells, generated *in vitro* or isolated from breast cancer patients, promoted the acquisition of mesenchymal-like breast cancer stemness properties. This cancer-stemness-inducing function was restricted to a myeloid subset expressing the glycoprotein CD52. Single cell transcriptomic- and surface proteome-based interactome analysis pointed towards membrane-bound TGF-β1 as a potential factor involved in cancer stemness induction. Functional inhibition of the TGF-β1 pathway blocked the emergence of cancer stem cells induced by suppressive myeloid cells. These results therefore identified the underlying mechanisms of a new tumor-promoting function of immunosuppressive myeloid cells, which may potentially be targeted.

**Highlights:**

- Immunosuppressive CD33^high^CD52^+^ myeloid cells induce mesenchymal-like cancer stem cells
- Cancer stemness induction requires membrane bound TGF-β1
- Blockade of the TGF-β1 pathway prevents cancer stemness induction

## Introduction

Escape from protective anti-tumor immunity is one of the hallmarks of cancers^1^. A primary strategy employed by tumors to avoid immune recognition and elimination consists in the induction of myeloid cells endowed with immunosuppressive activities^2^. Tumor-derived soluble and membrane-bound factors are responsible for the accumulation of these immunosuppressive myeloid cells in the primary tumor beds, lymphoid organs, bloodstream, and in the pre- and post-metastatic niches^3,4^. Suppressive myeloid cells encompass multiple phenotypically and functionally distinct subpopulations at different stages of differentiation, including tumor-associated neutrophils, tumor-associated macrophages or so-called Myeloid-Derived Suppressor Cells (MDSC)^5^. In many cancers, the accumulation of these immunosuppressive myeloid cell subsets has been correlated with disease progression, poor prognosis and resistance to therapies. In the specific case of breast cancers, considered for many years as immunologically “cold” due to a lack of infiltration with effector T lymphocytes, the tumor microenvironment contains a significant proportion of immunosuppressive cells of myeloid origin, which substantially contribute to disease development^6–8^. Suppressive myeloid cell elimination or inactivation conversely enhances responses to immunotherapeutic approaches^2,9–12^.

A common defining and unifying function of these myeloid cell subsets is their capability to inhibit anti-tumoral effector immune cells, including CD4^+^ T helper, CD8^+^ cytotoxic T lymphocytes and NK cells^3,4,13^. However, beyond this cardinal immunosuppressive property, these cells are also equipped with a broad spectrum of other tumor-promoting activities^14,15^. Indeed, different subpopulations of suppressive myeloid cells can directly enhance malignant cell survival and proliferation, contribute to tumor neo-angiogenesis and lymphangiogenesis, foster cancer cell invasion and metastasis, prepare the pre-metastatic sites to seeding by metastasizing neoplastic cells, and contribute to malignant cell resistance to radio- and chemotherapies^14^.

Some reports have suggested that immunosuppressive myeloid cells may promote the epithelial-to-mesenchymal transition (EMT) program in tumor, which might convey stem-like properties to cancer cells^16–24^. However, the impact of immunosuppressive myeloid cells in the acquisition and maintenance of cancer stemness has been explored in only a very limited number of studies^25–27^ and the underlying mechanisms are not fully comprehended. Cancer stem cells (CSC), a sparse subset of cancer cells defined by their unlimited ability to self-renew through asymmetrical division, are at the origin of tumor heterogeneity and are essential for tumor initiation, invasion, metastasis and resistance to therapies^28^. CSC thus represent a major cause of disease recurrence and patient relapse. A better understanding of the mechanisms by which these cells are maintained and enriched, and how they may potentially be eliminated is therefore of utmost importance.

In the present study, we demonstrated that human monocyte-derived suppressive cells generated *in vitro* or isolated from breast cancer patients were uniquely capable of inducing tumor cells exhibiting CSC properties. Suppressive myeloid cells primarily fostered the emergence of cancer cells associated with a mesenchymal-like CSC gene expression signature. A specific subset of suppressive myeloid cells, characterized by the expression of the glycoprotein CD52, was primarily responsible for cancer stemness induction. The combination of scRNAseq transcriptomic-, surface proteome (surfaceome)-based interactions studies and blockade experiments led to the identification of membrane TGF-β1 as a main factor involved in cancer stemness induction by suppressive myeloid cells.

## Results

### Myeloid-derived immunosuppressive cells generated *in vitro* or isolated from breast cancer patients induce the acquisition of cancer stemness properties

The potential contribution of immunosuppressive myeloid cells to cancer stemness induction was assessed using two complementary approaches. First, we took advantage of human monocyte-derived suppressor cells generated *in vitro* (HuMoSC)^29^ (**Figure S1A**). Because HuMoSC can be reproducibly generated in large number and because their immunosuppressive activity and development can be tightly controlled over time, this first approach allowed us to clearly analyze the cellular and molecular bases underlying their mode of action. The immunosuppressive function of HuMoSC, evidenced by their ability to suppress T lymphocyte proliferation was confirmed for each generated batch (**Figure 1A).** In a second approach, to validate the key findings obtained with HuMoSC, we analyzed suppressive myeloid cells isolated from breast cancer patients.

**Figure 1.**
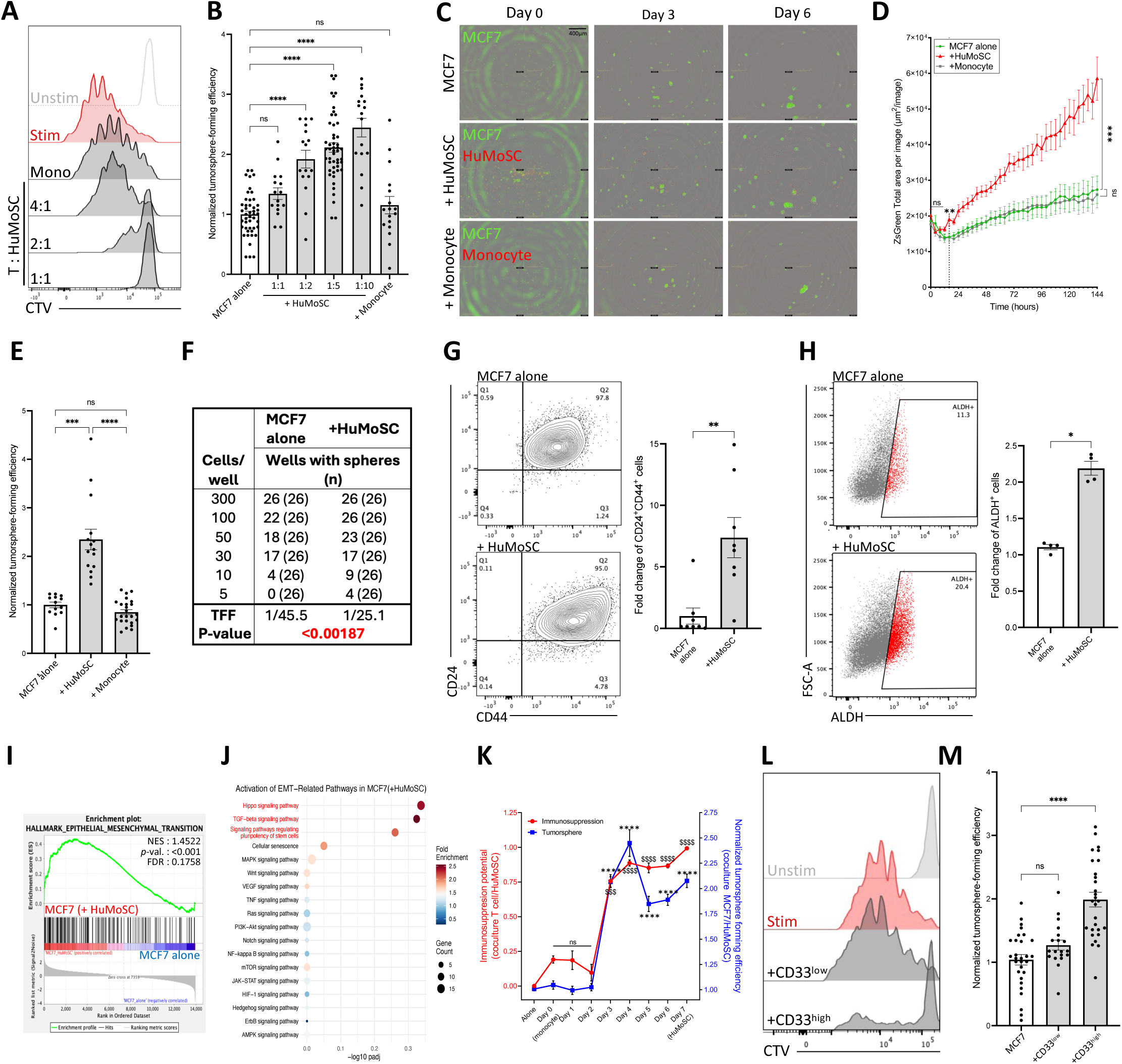
Immunosuppressive myeloid cells increase breast cancer stemness. (A) CellTrace Violet (CTV) dilution of unstimulated and CD3/CD28 stimulated T cells after 5-day co-culture with monocytes or HuMoSC at the indicated ratios (T cell:HuMoSC). (B) Tumorsphere formation assay of MCF7 cells alone or in co-culture with Human Monocyte-derived Suppressive Cells (HuMoSC) at the indicated ratio or monocytes at a 1:5 (MCF7:monocyte) ratio. (C) Live videomicroscopy representative images of ZsGreen-expressing MCF7 cells (green) alone or in co-culture with CellTrace Far Red-stained HuMoSC or monocytes (red) in tumorsphere forming conditions (MCF7:HuMoSC or monocyte ratio of 1:5). (D) Tumorsphere formation live videomicroscopy quantification of ZsGreen intensity over time of MCF7 cells alone (green) or in co-culture with HuMoSC (red) or monocyte (grey) (MCF7:HuMoSC or monocyte ratio of 1:5). Dashed line: 16h timepoint. (E-K) All analyses were performed on FACS-sorted MCF7 pre-cultured alone or with HuMoSC for 72 hours in adherent, classical 2D culture conditions (MCF7:HuMoSC or monocyte ratio of 1:5). (E) Normalized tumorsphere-forming efficiency of MCF7 cells pre-cultured alone or with HuMoSC or monocytes prior to seeding in tumorsphere forming conditions. (F) In vitro Extreme Limiting Dilution Assay (ELDA) of the tumorsphere formation ability of MCF7 cells. The number of wells presenting tumorspheres and total number of wells assayed in each condition is indicated (n). TFF, Tumorsphere Forming Frequency. (G and H) Representative FACS plots (left) and normalized proportion (right) of CD44^+^CD24^-^ MCF7 cells (G) and ALDH^+^ MCF7 cells (H). (I) GSEA plot showing the Normalized Enrichment Score (NES) and associated p-value for the EMT Hallmark gene set in the variance-ranked genes for MCF7 cells cultured 24h with HuMoSC compared to MCF7 cells cultured alone. FDR: False Discovery Rate. (J) Bubble plot showing the upregulation of EMT-related Kegg signaling pathways in MCF7 cultured for 24 hours with HuMoSC compared to MCF7 cultured alone. Pathways of interest are highlighted in red. (K) Monocyte-to-HuMoSC immunosuppression potential upon T-cell coculture (left, red) and MCF7 tumorsphere formation assay after co-culture with HuMoSC (right, blue) at each day of their polarization from monocytes (Day 0) to HuMoSC (Day 7). (L) CellTrace Violet dilution of T cells unstimulated (Unstim), stimulated with anti-CD3/CD28 (Stim) or stimulated and co-cultured with CD33^low^ or CD33^high^ myeloid cells isolated from pleural effusion of breast cancer patients. (M) Tumorsphere formation assay of MCF7 cells alone or in co-culture with CD33^low^ or CD33^high^ isolated from pleural effusion of breast cancer patients. Data are means ± S.E.M. ∗p < 0.05, ∗∗p < 0.01, $$$/∗∗∗p < 0.001, $$$$/∗∗∗∗p < 0.0001. ns, not significant. Kruskal-Wallis test with Dunn’s multiple comparisons post-test (B, D, E, K, M); Tumorsphere Forming Frequency (TFF) and associated p-value was calculated using ELDA software (F); Student’s unpaired t-test (G, H).

The quantification of cancer cells exhibiting stemness properties conventionally relies on tumorsphere-forming assays. In the selective culture conditions required for these tests, only a sparse subpopulation of cells exhibiting stemness activities are able to survive and self-renew. We determined that HuMoSC significantly enhanced the formation of MCF7 breast cancer cell tumorspheres in a ratio-dependent manner (**Figure 1B and Figure S1B**). Importantly, the monocytes from which HuMoSC were generated, were not immunosuppressive (**Figure 1A**) and had no significant effect on tumorsphere formation (**Figure 1B**). The kinetic of tumorsphere formation and the interactions between HuMoSC and cancer cells in CSC-like-selecting conditions were then analyzed by time-lapse videomicroscopy. The results depicted in **Supplemental Video 1-3** and **Figure 1C and D** indicated that ZsGreen-expressing MCF7 tumorsphere formation increased over time upon co-culture at a 1:5 ratio with Cell Trace Red-stained HuMoSC, but not with Cell Trace Red-stained monocytes. Tumorsphere promotion by HuMoSC was significantly detectable as early as 16 hours after the initiation of the co-culture (**Figure 1D**). These results thus suggest that only immunosuppressive myeloid cells can foster breast cancer cell stemness properties.

To evaluate the stability of stemness induction, MCF7 cells co-cultured with HuMoSC for 72 hours in 2D conditions were purified by FACS (CD45-based HuMoSC elimination) and incubated in tumorsphere-forming conditions without HuMoSC. A significant increase in tumorsphere formation was again observed when MCF7 were pre-cultured with HuMoSC compared to MCF7 cultured alone or pre-cultured with non-immunosuppressive monocytes (**Figure 1E**). This result suggests that HuMoSCs induce lasting cancer stemness properties in MCF7 tumor cells. *In vitro* extreme-limiting dilution assays further indicated that HuMoSC promoted a significant increase in the frequency of cells with CSC functional properties in the pool of bulk cancer cells (**Figure 1F**). Consistent with these functional results, flow cytometry analysis indicated that a significant increase in both CD44^+^CD24^-^ cells and ALDH positive cells was detected after co-culture of MCF7 tumor cells with HuMoSC (**Figure 1G and 1H**).

EMT is a critical and dynamic process associated with CSC induction. The EMT program depends on signalling pathways such as TGF-β, Wnt/β-catenin, Notch and Hedgehog, leading to the upregulation of EMT-associated transcription factors^30^. To explore the impact of HuMoSC on MCF7 EMT-associated signalling pathways, we performed bulk RNA barcoding (BRB) sequencing analyses of MCF7 cells purified after 24 hours post co-culture with HuMoSC, compared to MCF7 cultured alone. Consistently, gene set enrichment analysis (GSEA) demonstrated an enrichment for the MSigDB Hallmark EMT gene set in MCF7 co-culture with HuMoSC (**Figure 1I**). Interestingly, Kegg signaling pathway enrichment analysis revealed an upregulation of the EMT-controlling Hippo and TGF-β signaling pathways in MCF7 co-cultured with HuMoSC (**Figure 1J**). Likewise, signalling pathways involved in stem cell pluripotency, an established feature of CSC^31^, were upregulated by HuMoSC in MCF7 (**Fig 1J**). Altogether, these transcriptomic results indicate that HuMoSC induce the expression of genes and pathways related to EMT and stemness in MCF7 cancer cells.

We then analyzed the kinetics of acquisition of the immunosuppressive function and cancer stemness-promoting capability of the cells during monocyte (Day 0) differentiation towards HuMoSC (Day 7) (**Figure S1A**). We demonstrated that myeloid cells concomitantly acquired suppressive activities and tumorsphere-promoting properties, between day 2 and day 3 in the differentiation process from monocytes to HuMoSC (**Figure 1K and S1D-E)**.

In a second approach, to extend these findings to a clinical setting, myeloid cells were isolated from pleural effusions or ascites of breast cancer patients based on the expression of CD33, a member of the sialic acid-binding immunoglobulin-like lectin (Siglec) family expressed on myeloid cells^32^. Two distinct populations of myeloid cells differentially expressing CD33 (CD33^low^ and CD33^high^) were identified (**Figure S1C**). These cells have been associated with granulocytic-like and monocytic-like myeloid cells, respectively^33^. CD33^high^ myeloid cells were more immunosuppressive than their CD33^low^ counterparts (**Figure 1L)**. Importantly, immunosuppressive CD33^high^ but not non-immunosuppressive CD33^low^ myeloid cells were capable of promoting MCF7 cells tumorsphere formation (**Figure 1M**).

Altogether, these results indicate that immunosuppressive myeloid cells generated *in vitro* or isolated from patients, but not their non-immunosuppressive counterparts, foster the acquisition of cancer stemness properties.

### Human monocyte-derived immunosuppressive cells induce mesenchymal-like cancer stem cells

Liu *et al.* have identified distinct subsets of breast CSC exhibiting plasticity towards epithelial (E-CSC) or mesenchymal (M-CSC) characteristics^34^. While E-CSC seem to be located in the inner core of the tumor and exhibit a proliferative phenotype, their mesenchymal counterparts are located at the invasive edge of the tumor and have been shown to exhibit high invasive properties^34,35^. To precisely investigate the impact of HuMoSC on CSC heterogeneity and plasticity on a per cell basis, we performed single cell RNA-sequencing (scRNAseq) analyses of 1) MCF7 or HuMoSC cultured alone in conventional culture conditions “2D-Culture”, allowing for the separate analysis of gene expression profile by cancer cells or myeloid cells in non-selective conditions; 2) MCF7 or HuMoSC cultured separately for 3 days in tumorsphere-forming conditions “Tumorsphere conditions”; and 3) “Tumorsphere co-cultures”: MCF7 cultured for 3 days with HuMoSC in tumorsphere-forming conditions, as described in **Figure 2A**. Following exclusion of stressed^36^ and dead cells and putative doublet removal (**Figure S2A**), we analyzed the transcriptome of 8.643 HuMoSC (∼3.200 genes/cell) and 7.259 MCF7 (∼5.300 genes/cell) across all conditions. Following data integration, four Louvain clusters of MCF7 cells, defined as *PTPRC*^-^ (CD45^-^), and four Louvain clusters of HuMoSC (*PTPRC^+^*) were identified (**Figure 2B and Figure S2B**).

**Figure 2.**
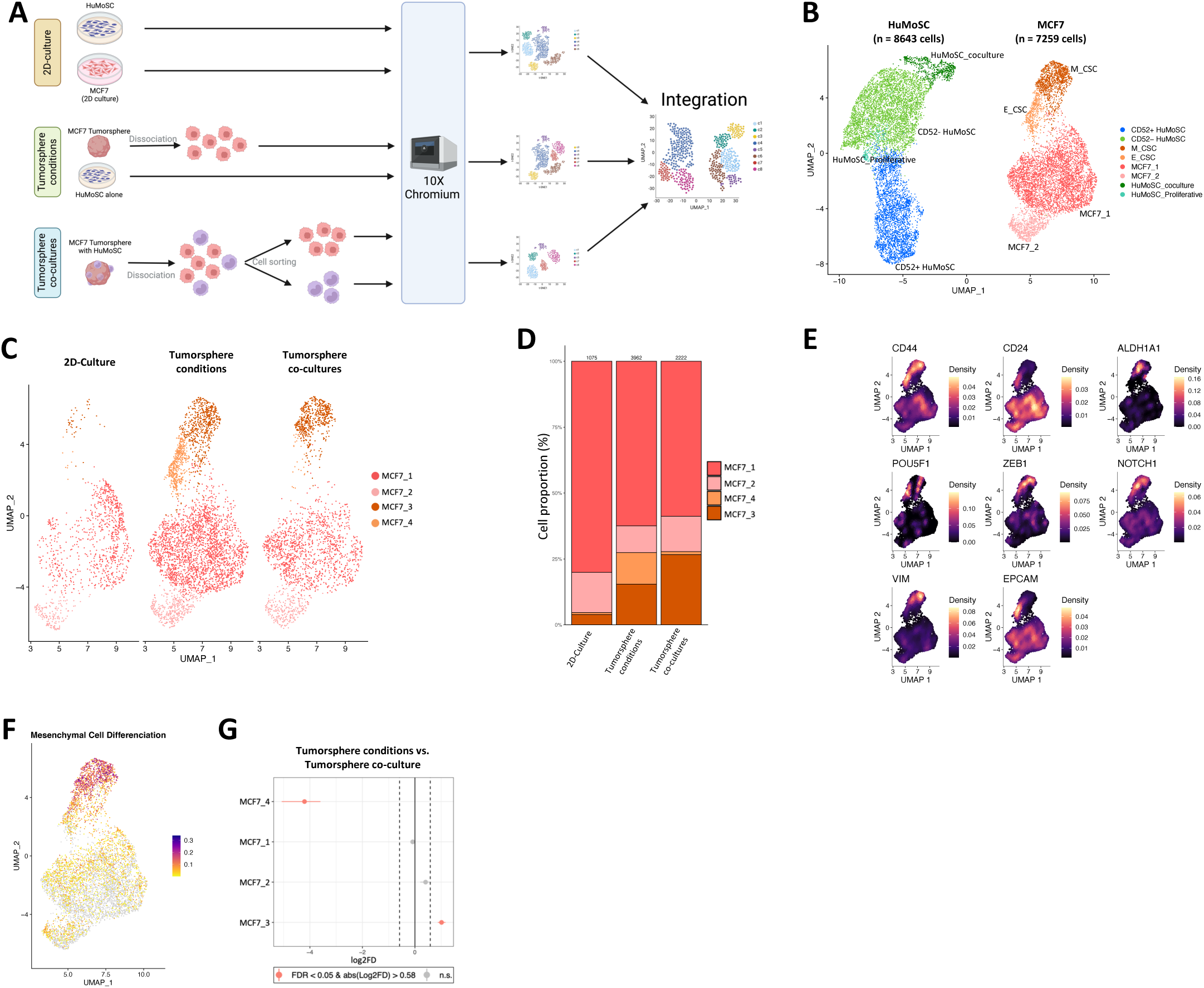
HuMoSC foster Mesenchymal-like Cancer Stem Cell emergence. (A) scRNAseq workflow of the three different conditions. See also Figure S2A. (B) Two-dimensional UMAP of scRNA-seq data of HuMoSC (n = 8643 cells) and MCF7 (n = 7259 cells) after integration of the three experimental conditions “2D-culture”, “Tumorsphere conditions” and “Tumorsphere co-cultures”. (C) Split two-dimensional UMAP of MCF7 cells based the three experimental conditions. (D) Bar plot of cell proportion of each MCF7 cluster in each condition. (E) Density Plot representing the expression of CSC and EMT genes in MCF7 cell clusters. (F) Feature Plot representation of core signature associated with the 55 genes from Mesenchymal Cell Differentiation gene set. (G) Relative differences in cell proportions for each MCF7 cluster between the “Tumorsphere conditions” versus “Tumorsphere co-culture” conditions.

We first focused on precisely annotating the four clusters of MCF7 cells and on determining whether CSC (enriched in the tumorsphere forming conditions) could be distinguished from non-CSC (predominant in the “2D-Control” conditions). The comparison of the four clusters of MCF7 in the different experimental conditions indicated that the clusters “MCF7_3” and “MCF7_4” were present almost exclusively in the “Tumorsphere conditions” (in culture conditions promoting the survival of cancer cells with stemness properties), while the “2D-Culture” condition is mainly composed of the clusters “MCF7_1” and “MCF7_2” (**Figure 2C and 2D**). This observation suggested that “MCF7_1” and “MCF7_2” clusters correspond to “non-CSC-like cells” while the “MCF7_3” and “MCF7_4” clusters relate to CSC-like cells. To confirm this hypothesis, we next analysed the expression of genes allowing for the identification of breast cancer CSC. Consistent with our initial analysis (**Figure 1G-H)**, “MCF7_3” and “MCF7_4” clusters strongly expressed *ALDH1A1* and *CD44* but had lost the expression of *CD24* (**Figure 2E**). Moreover, these two clusters expressed genes associated with the cancer stemness-associated EMT program such as *POU5F1* (encoding for OCT4), *ZEB1*, *NOTCH1* and *VIM* (**Figure 2E**). Altogether, these data pinpointed “MCF7_3” and “MCF7_4” clusters as CSC-like cells.

We next sought to specifically investigate the differences between the two non-CSC-like clusters (“MCF7_1” and “MCF7_2”) and the two CSC-like clusters (“MCF7_3” and “MCF7_4”). Cell cycle scoring (**Figure S2C**) and expression of proliferation-associated gene (*MKI67*; *STMN1* **Figure S2D**) analyses indicated that the two distinct clusters of non-CSC could be distinguished based on their proliferating states. Deeper analyses of the CSC-like clusters highlighted a strong expression of the epithelial marker *EPCAM* in “MCF7_4” (**Figure 2E**). We thus hypothesized that the two CSC-like clusters might represent phenotypically distinct CSC, harbouring an epithelial or a mesenchymal phenotype. To investigate the mesenchymal phenotype of the “MCF7_3” cluster, we used a scoring method based on the “Mesenchymal Cell Differentiation” gene set extracted from Harmonizome 3.0^37^, a gene signature encompassing 55 genes associated with cellular acquisition of mesenchymal features (**Figure 2F**). This gene signature specifically discriminated “MCF7_3” cluster from the other clusters, thereby confirming the mesenchymal phenotype of these cells. Interestingly, only the mesenchymal CSC (“MCF7_3”) cluster and not the epithelial CSC (“MCF7_4”) cluster was detected in MCF7-HuMoSC co-cultures (“Tumorsphere co-cultures” conditions, **Figure 2C, right panel and Figure 2D**). Strengthening this observation, a statistical enrichment test validated the enrichment of cancer cells with the mesenchymal “MCF7_3” cluster upon co-culture with HuMoSC (**Figure 2G**). These results strongly suggest that HuMoSC may promote CSC plasticity towards mesenchymal CSC.

Altogether, results from these scRNAseq analyses discriminated four clusters of MCF7 cells, two of which are non-CSC at different phases of the cell cycle (“MCF7_1” and “MCF7_2” clusters) and two clusters of CSC, characterized by mesenchymal (“MCF7_3”) or epithelial (“MCF7_4”) phenotypes. Based on these data, for better clarity in further analyses, we renamed these two CSC clusters “M_CSC” (for mesenchymal, MCF7_3) and “E_CSC” (for epithelial, MCF7_4), respectively (**Figure 2B**). Immunosuppressive myeloid cells foster the emergence of M_CSC, previously characterized by a highly invasive phenotype^34^.

### A specific subset of immunosuppressive myeloid cells expressing CD52 is capable of inducing cancer stemness

HuMoSC, like myeloid suppressive cells described in cancer patients, are constituted of different subsets. To determine whether dedicated subsets within the HuMoSC population may be specifically endowed with cancer stemness-promoting functions, scRNAseq analyses of HuMoSC in the 3 different experimental conditions described hereabove (**Figure 2A**, **2B and Figure S2A)** were initially performed. Four distinct HuMoSC subsets were identified across all integrated conditions (**Figure 2B**). When analysing the distribution of the cells in each cluster in the different experimental conditions (**Figure 3A and 3B**), we identified two core populations of HuMoSC present in each condition (“HuMoSC_1” and “HuMoSC_2). We also identified one cluster characterized by its proliferative profile (“HuMoSC_4”), as depicted by cell cycle scoring and the expression of cell cycle markers (*MKI67* and *STMN1)* (**Figure S2C and S2D**). We further annotated this cluster as “HuMoSC_Proliferative”. Interestingly, the cluster “HuMoSC_3” only appeared in MCF7-HuMoSC co-cultures (“Tumorsphere co-cultures”). Enrichment analysis statistically confirmed that this “HuMoSC_3” cluster specifically arose in the co-culture with MCF7 cancer cells in selective tumorsphere-forming conditions (**Figure 3C**). For better clarity, we thus annotated the “HuMoSC_3” cluster as “HuMoSC_coculture” for further analysis. Of note, the “HuMoSC Proliferative” cluster was significantly depleted in this co-culture condition. We next sought to analyse the differentiation potency of these 4 HuMoSC clusters by performing a cytoTRACE-based transcriptional analysis of HuMoSC differentiation status. CytoTRACE assigns a score to each cell based on the diversity of expressed genes, which serves as a proxy for the cell differentiation potential. Cells with higher CytoTRACE scores exhibit broader gene expression profiles and are predicted to be less differentiated, indicating higher potency and stem-like characteristics. Conversely, cells with lower scores are more differentiated and exhibit more specialized gene expression profiles^38^. This method predicts the differentiation potential of clusters based on the expression level of specific genes associated with specific developmental states^38^. Based on cytoTRACE predicted ordering, the “HuMoSC_coculture” cluster exhibited the most differentiated state, while “HuMoSC_2” demonstrated the highest potency and thus the least differentiated state (**Figure 3D**). MDSC exhibit an “immature” and less differentiated phenotype compared to other myeloid cells^39^. In addition to the CytoTRACE analysis, we applied an MDSC gene signature obtained from PangloDB database to our single cell dataset (**Figure 3E)**. The less differentiated clusters predicted with CytoTRACE (“HuMoSC_1” and “HuMoSC_2”) exhibited a strong positive correlation with the MDSC gene signature while the “HuMoSC_coculture” cluster lacked this signature (**Figure 3E**). Considering the “immature” nature of MDSC^39^, this result is coherent with the low differentiation potential of “HuMoSC_1” and “HuMoSC_2” clusters predicted with the cytoTRACE analysis.

**Figure 3.**
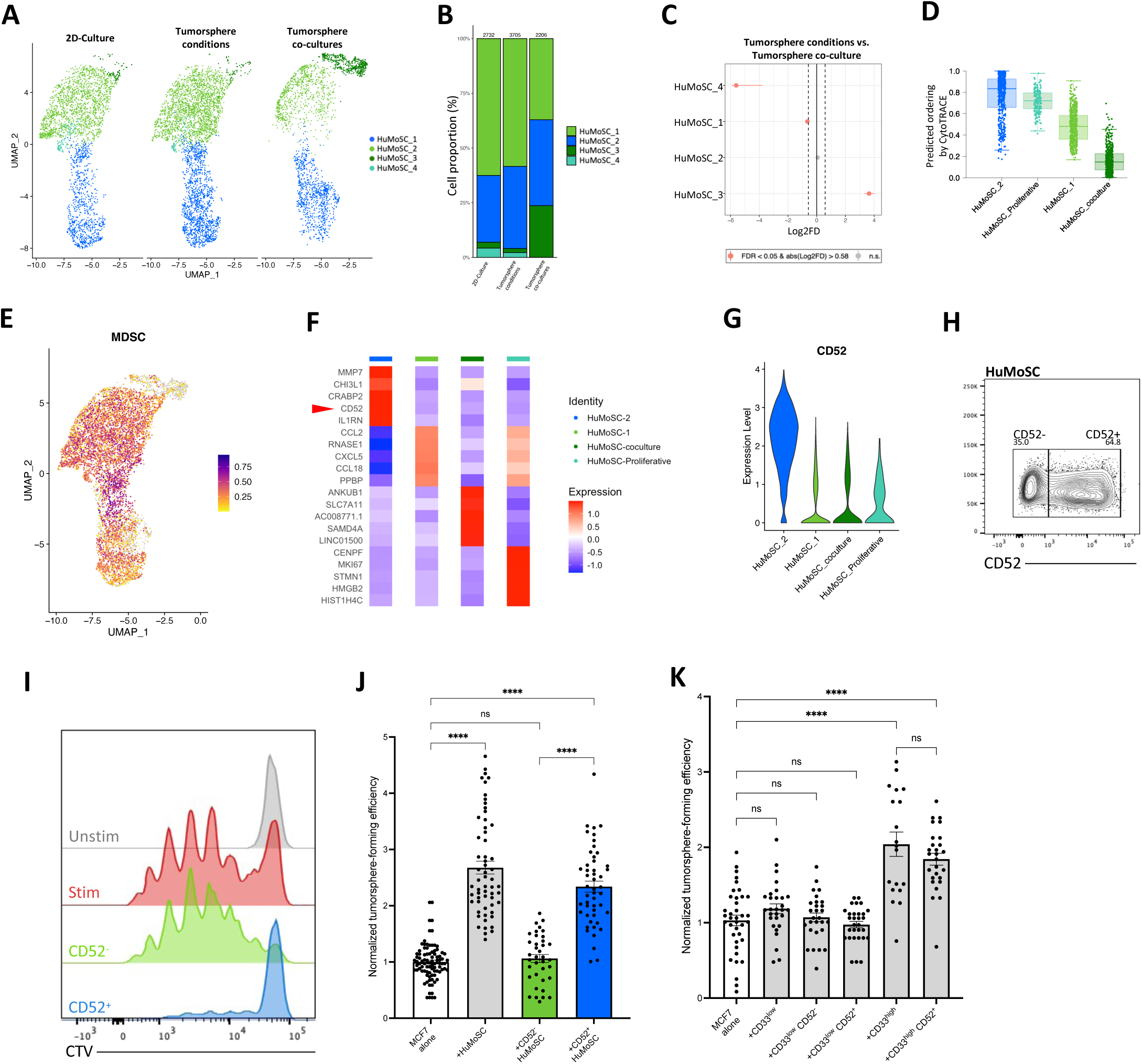
CD52^+^ HuMoSC are immunosuppressive and stemness-inducting cells. (A) Split two-dimensional UMAP of HuMoSC cells based the three experimental conditions. (B) Bar plot of cell proportion of each HuMoSC cluster in each condition. (C) Relative differences in cell proportions for each HuMoSC clusters between “Tumorsphere conditions” and “Tumorsphere co-culture” conditions. (D) Boxplot showing the predicted ordering by cytoTRACE as assessment of cells differentiation potency. (E) Feature Plots representation of core signature associated with “MDSC” gene sets extracted from PangloDB. (F) Top 5 most significant differentially expressed genes across all HuMoSC clusters. (G) Violin plot expression of CD52 gene across HuMoSC clusters. (H) CD52 flow cytometry analysis of HuMoSC. See also Figure S3A. (I-J) CellTrace Violet dilution of T cells (I) and MCF7 tumorsphere formation assay (J) after co-culture with total, CD52^-^ and CD52^+^ HuMoSC at a 1:5 ratio (MCF7:HuMoSC). (K) Tumorsphere formation assay of MCF7 cells alone or in co-culture with myeloid cells isolated from pleural effusion of breast cancer patients and differentially expressing CD33 (high or low) and CD52. See also Figure S3B. (J and K) Statistical significance was assessed using a Kruskal-Wallis test with Dunn’s multiple comparisons test. ****p<0.0001; ns, not significant. Data are represented as mean values ± S.E.M.

In an effort to further identify cell surface markers that would allow subset discrimination, a Differential Gene Expression (DGE) analysis was performed. CD52 was identified as the most discriminating cell surface marker between “HuMoSC_1” and “HuMoSC_2” (**Figure 3F and Figure 3G**). CD52 is a 12 amino-acid glycoprotein expressed by various immune cells, including T and B lymphocytes, monocytes, and dendritic cells, and playing a role in modulating immune responses^40^. Consistent with the scRNAseq data, flow cytometry analyses further indicated that two distinct populations could be discriminated within total HuMoSC based on CD52 expression (**Figure 3H**). Of note, human monocytes expressed CD52 homogeneously as previously reported^41^ (**Figure S3A**). Based on this observation, we then re-named the clusters HuMoSC_1 and HuMoSC_2 as “CD52^-^ HuMoSC” and “CD52^+^ HuMoSC”, respectively (**Figure 2B**).

Based on these data, we next purified CD52^-^ and CD52^+^ HuMoSC by FACS and analysed both their immunosuppressive potential and their cancer stemness promoting capability. Interestingly, only CD52^+^ HuMoSC were endowed with both immunosuppressive functions and the ability to significantly increase MCF7 tumorsphere formation, while CD52^-^ HuMoSC were neither immunosuppressive nor capable of triggering cancer stemness (**Figure 3I and 3J**). We next interrogated the value of CD52 to discriminate specialized myeloid cell subsets in pleural effusions from breast cancer patients. Only CD33^low^ cells differentially expressed CD52, while all CD33^high^ cells appeared to co-express CD52 (**Figure S3B**). Similar to the results obtained with total CD33^high^ cells (**Figure 1L**), only purified CD33^high^CD52^+^ cells were able to promote CSC-like cell emergence, while CD33^low^ cells did not significantly increase tumorsphere formation, independently of their expression of CD52 **(Figure 3K)**.

Altogether, these results uncover that a dedicated CD52-expressing subset of suppressive myeloid cells is primarily responsible for cancer stemness promotion.

### Suppressive myeloid cells induction of cancer stemness depends on membrane-bound factors

We next focused on identifying the mechanisms underlying cancer stemness induction by suppressive myeloid cells. First, the nature of the cellular interactions between HuMoSC and MCF7 cells were analyzed. Quantification of the overlaying signals from HuMoSC (red) and MCF7 tumorspheres (green) in time-lapse videomicroscopy experiments indicated a close contact between the two cell types, with maximum interactions detected after 24 hours of co-culture in CSC-enriched conditions. Conversely, monocytes interacted with MCF7 to a much lower extend compared to HuMoSC (**Figure S4A and Supplemental Video 1 to 3**). Live immunofluorescence analysis further confirmed the presence of HuMoSC in direct contact with tumor cells (**Figure 4A**). Based on these observations, we next investigated the possible involvement of soluble and membrane-bound factors in the induction of CSC by immunosuppressive myeloid cells. The physical separation of HuMoSC from tumor cells by a transwell insert abrogated the effects of these suppressive myeloid cells on tumorsphere formation, indicating that HuMoSC-mediated promotion of cancer stemness depends on a cell-to-cell contact (**Figure 4B**). Furthermore, neither HuMoSC-conditioned medium (CM_HuMoSC_) nor the supernatant of HuMoSC-MCF7 co-cultures (CM_co-culture_) increased tumorsphere formation **(Figure 4B)**. These results strongly suggest, first that the mechanism of cancer stemness induction by HuMoSC did not directly depend on secreted active factors but rather requires a direct cell-to-cell contact, and second that HuMoSC-MCF7 interaction did not lead to the secretion of products directly enhancing cancer stemness.

**Figure 4.**
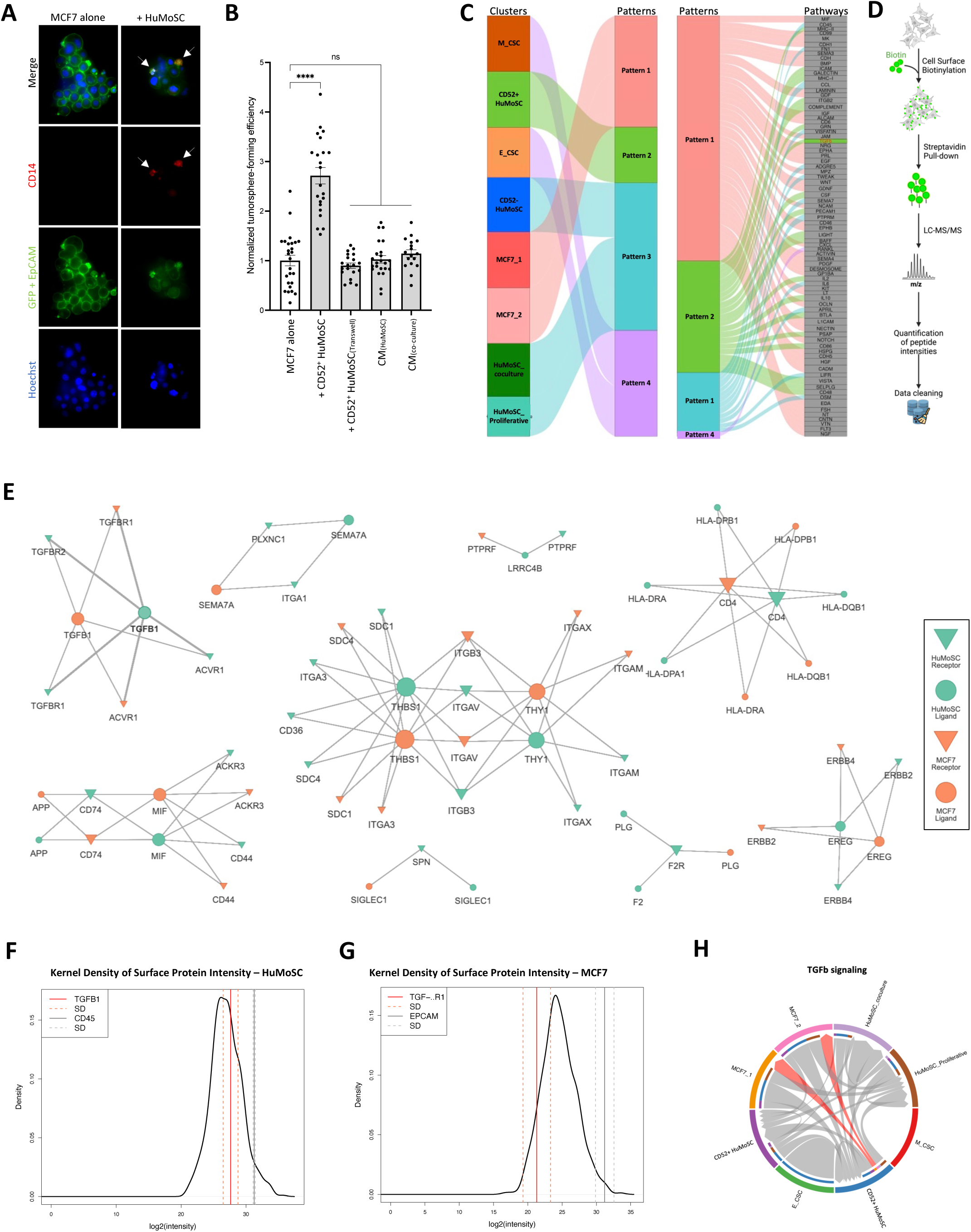
Stemness promotion requires a direct cell-to-cell contact with transcriptomic and proteomic prediction of TGF-β involvement. (A) Immunofluorescence images of live ZsGreen-expressing MCF7 tumorspheres alone or in co-culture with HuMoSC (1:5 ratio) after staining with anti-CD14 (red) anti-Epcam (green) antibodies. Nuclei stained with Hoechst (blue). All images are representative of at least three replicates. White arrows points towards tumorspheres-associated HuMoSC. (B) Tumorsphere formation assay of MCF7 cells cultured alone, with CD52^+^ HuMoSC or with conditioned medium (CM) from HuMoSC (CM_HuMoSC_) or from co-culture of MCF7 and HuMoSC (CM_co-culture_). In some conditions, MCF7 were physically separated from HuMoSC cells by a transwell. Statistical significance was assessed using a Kruskal-Wallis test with Dunn’s multiple comparisons test. ***p<0.001; ns, not significant. Data are represented as mean values ± S.E.M. (C) CellChat inferred outgoing communication patterns of the different cell clusters (left panel) and associated signaling pathways (right panel). (D) Surfaceome workflow for the identification and quantification of total cell surface proteins. (E) Talkien-predicted protein interaction network of HuMoSC and MCF7 surfaceome. (F-G) Kernel density plot showing log2 intensity of TGFB1 in HuMoSC surfaceome (F) and TGF-BR1 (G) in MCF7 surfaceome. CD45 (F) and EPCAM (G) were used as reference for protein intensities. Dashed line: calculated standard deviation (SD) of the three replicates for the corresponding protein. (H) Chord diagram showing predicted TGFb signaling between clusters. Red arrows highlight the signaling specificity of CD52^+^ HuMoSC towards MCF7_1 and MCF7_2 clusters.

### Combining transcriptome and surfaceome analysis identified TGF-β1 as a key factor involved in HuMoSC and MCF7 interactions

The observation that cancer stemness induction by HuMoSC depends on a direct cellular contact implies the engagement between surface molecules expressed by both cell types, triggering signalings responsible for the activation of the cancer stemness program in MCF7 tumor cells. To identify these putative interacting molecular candidates, we employed a multi-omic strategy that integrated cell-cell communication predictions from our scRNAseq with a comprehensive cell surface proteome analysis. This transcriptomic and proteomic combinatorial approach allowed us to establish a restricted predictive list of interacting candidates between HuMoSC and MCF7 cells for further functional investigations.

CellChat analysis allows for the inference of cell-to-cell communication using scRNAseq data, by applying a method of pattern recognition using non-negative matrix factorization to uncover both the overarching communication patterns and crucial signals within distinct cell clusters^42^. We established a global communication analysis of the 8 clusters from our scRNAseq, combining the HuMoSC and MCF7 analysis. First, we identified the dominant outgoing cell communication patterns of each cluster, allowing for a similarity categorization of cell signalings (**Figure 4C**). This analysis is informative to unify the different clusters of a complete dataset based on their similar capacity to induce a signaling to any potential receiver cell. This predictive analysis highlighted four coherent patterns. Pattern 1 and pattern 4 regrouped the outgoing signaling pathways associated with non-CSC (“MCF7_1” and “MCF7_2”) and CSC (“M_CSC” and “E_CSC”), respectively. Interestingly, the immunosuppressive, stemness-inducing “CD52^+^ HuMoSC” cluster was part of a unique pattern of its own (Pattern 2) and all other HuMoSC clusters were part of a same pattern (Pattern 3) (**Figure 4C**). This “CD52^+^ HuMoSC”-specific outgoing communication pattern 2 was associated with twenty signaling networks (*MHC-II*, *ICAM*, *GALECTIN*, *MHC-I*, *ITGB2*, *COMPLEMENT*, *GRN*, *TGFb*, *CSF*, *PECAM1*, *LIGHT*, *BAFF*, *IL2*, *IL-10*, *BTLA*, *PSAP*, *CD86*, *VISTA*, *SELPLG* and *CD48*).

Because this CellChat predictive interaction analysis remained limited to transcriptomic data, and to narrow down the number of potential interacting cell surface candidate molecules, we ran an exhaustive cell surface proteome (Surfaceome) investigation of MCF7 and HuMoSC^43^. As depicted in **Figure 4D**, total surface proteins of live HuMoSC or MCF7 cells were separately biotinylated and after streptavidin-based affinity purification, total surface proteins were analyzed by Liquid Chromatography coupled to tandem Mass Spectrometry (LC-MS/MS). We cleaned the data based on the non-null quantification of each triplicate and excluded from the analysis the proteins identified with a SPAT score < 8, as previously reported^44^. This SPAT scoring method allows to grade proteins according to their cell surface location probability, based on a unified annotation database. A total of 611 surface proteins were identified on HuMoSC and 488 on MCF7 (**Supplemental Table 1**). We next created a predictive proteomic-based interaction network between HuMoSC and cancer cells using the TALKIEN algorithm after specification of using the CellChat Database only to compare the proteomic prediction with the transcriptomic prediction (**Figure 4E**). Among the predicted interaction networks, only *TGFβ* and *MHC-II* signaling pathways aligned with the twenty predicted candidates from the CellChat transcriptomic analysis. TGF-β1 is described as a major actor of EMT initiation and stemness induction^45,46^. However, TGF-β1 has widely been reported to function as a soluble cytokine, thus classifying the TGF-β1 signaling as a “secreted pathway” and not as a “cell-to-cell contact” communication pathway in the CellChat database. Because our data pointed toward a direct cell-cell contact-dependent mechanism of cancer stemness induction (**Figure 4B**), we sought to further sort out the role of the TGF-β1-related communication pathway by analyzing the expression of TGF-β1, TGF-βR1 and ACVR1 (actors in the TGF-β-dependent interaction network as shown in **Figure 4E**) in Kernel density plots. TGF-β1 was present in high quantity in the pool of analyzed HuMoSC surface proteins (**Figure 4F**) and its binding partner receptors, TGF-βR1 (**Figure 4G**) and ACVR1 (**Figure S4B**) were detected in the membrane fraction of MCF7 proteins. In light of these last results, TGF-β1-centered CellChat re-analysis of the scRNAseq data highlighted that only “CD52^+^ HuMoSC”, but not “CD52^-^ HuMoSC”, displayed TGF-β-associated signaling communications towards the MCF7_1 and MCF7_2 clusters (**Figure 4H**).

Thus, using this innovative approach combining transcriptomics and proteomics analysis, we identified a potential candidate involved in the direct cell-to-cell, stemness-inducing interactions between HuMoSC and MCF7. Our analyses indeed revealed that TGF-β1 was present in the pool of HuMoSC total surface proteins, and the TGF-β pathway was predicted as a potent outgoing communication signaling from only immunosuppressive, cancer stemness-inducing “CD52^+^ HuMoSC” towards cancer cells.

### Membrane-bound TGF-β1 is required for cancer stemness induction by immunosuppressive myeloid cells

The possibility that TGF-β1 functions as a membrane-bound protein has only been documented in few reports, primarily related to regulatory T cells^47–53^ and its role in myeloid cells remains to be defined. We first determined that TGF-β1 secretion was not increased over time in HuMoSC culture supernatants compared to the supernatant of monocytes which were not able to induce cancer stemness (**Figure 5A**). Likewise, the co-culture of MCF7 and HuMoSC in direct contact did not increase TGF-β1 secretion (**Figure 5B**). Conversely, comparative surfaceome analysis of HuMoSC and monocytes from which HuMoSC were generated indicated that 73 cell membrane proteins were significantly overexpressed (**Supplemental Table 2 and 3**), among which TGFΒ1, and 129 significantly downregulated in HuMoSC compared to monocytes (**Figure 5C**). Consistently, western blot analysis of TGF-β expression displayed a strong overexpression in the pulled-down membrane protein fraction of HuMoSC compared to that of monocytes (**Figure 5D**).

**Figure 5.**
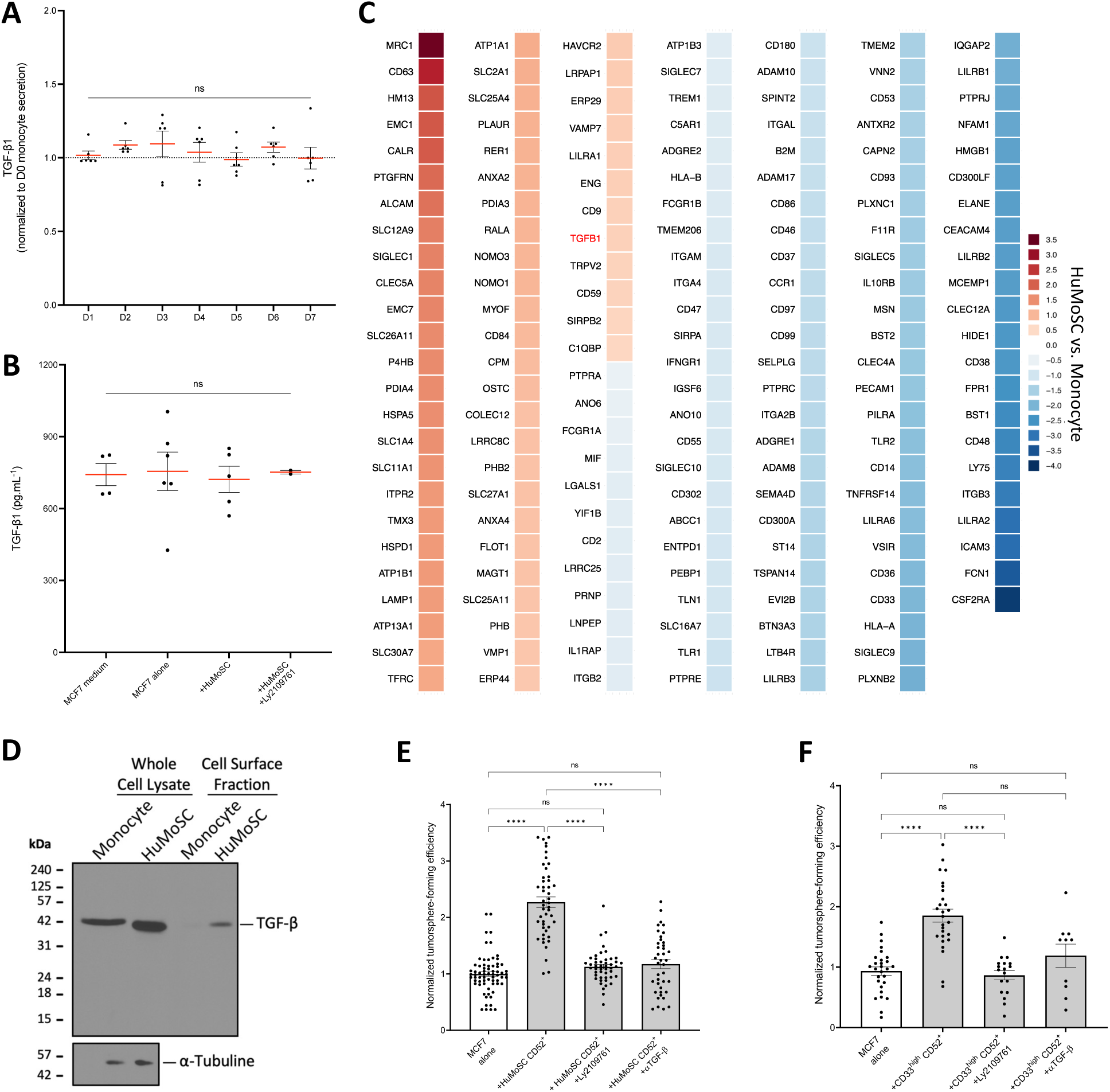
Membrane TGF-β1 and its associated pathway are involved in cancer stemness induction by suppressive myeloid cells. (A and B) Secreted TGF-β1 quantification by ELISA assay on supernatants from D0 to D7 HuMoSC (A) and supernatant from MCF7 cells alone or in co-culture with HuMoSC, with or without treatment with Ly2109761 (B). Secretion was normalized to D0-monocyte secretion (A). “MCF7 medium” condition correspond to basal TGF-β detected in the medium (B). (C) Heatmap comparing all significant surfaceome changes in HuMoSC compared to monocyte. (D) TGF-β western blot using Monocyte or HuMoSC total protein fractions before streptavidin pulldown (Whole Cell Lysate) or cell surface protein fractions of biotinylated proteins only (Cell Surface Fraction). α-Tubuline was used as a control for whole cell lysate loading. (E) Tumorsphere formation assay of MCF7 cells alone or after co-culture with CD52^+^ HuMoSC and treated or not with 5 μM TGF-βRI/II inhibitor Ly2109761 or 1 μg/mL αTGF-β neutralizing antibody. (F) Tumorsphere formation assay of MCF7 cells alone after in co-culture with CD33^high^CD52^+^myeloid cells isolated from pleural effusion of breast cancer patients and differentially expressing CD33 (high or low) and CD52 and treated or not with 5 μM TGF-βRI/II inhibitor Ly2109761 or 1 μg/mL αTGF-β neutralizing antibody. Data are means ± S.E.M. ∗p < 0.05, ∗∗p < 0.01, ∗∗∗p < 0.001, ∗∗∗∗p < 0.0001. ns, not significant. Kruskal-Wallis test with Dunn’s multiple comparisons post-test (A-B, E-F). Data are means of n = 3 independent biological replicates. (C) Log2 FC (HuMoSC/Monocyte) above 2 or below -2 (4-fold) were considered as significantly upregulated (red) or downregulated (blue), respectively. P-values were determined by unpaired two-tailed Student’s t-tests; P < 0.05.

Based on these data, and to further confirm the predicted role of TGF-β signaling in HuMoSC-induced cancer stemness, we next assessed whether blockade of different elements of this signaling pathway affected HuMoSC induction of cancer stemness. Our results indicate that while MCF7 treatment with TGF-β1 effectively increase MCF7 tumorsphere formation (**Figure S5A**), inhibition of TGF-βRI/II with the small molecule inhibitor Ly2109761^54^ abrogated the induction of tumorspheres by CD52^+^ HuMoSC (**Figure 5E**). Of note, no direct effect of the Ly2109761 inhibitor on MCF7 tumorsphere formation alone was observed, confirming that the observed inhibition is dependent on the co-culture settings (**Figure S5A**). Consistent with these data, a TGF-β1,2,3 capture antibody added in HuMoSC and MCF7 co-cultures also led to the inhibition of HuMoSC-induced tumorsphere formation (**Figure 5E**), and successfully blocked the tumorsphere-promoting effect of TGF-β1 on MCF7 (**Figure S5A**). Of clinical relevance, we lastly demonstrated that both inhibitors significantly inhibited tumorsphere promotion by CD33^high^CD52^+^ immunosuppressive cells from cancer patients (**Figure 5F**). Consistent with the literature^55^, this last result suggests that a similar signaling pathway involving TGF-β signaling contributes to cancer stemness promotion by patient-derived immunosuppressive myeloid cells. Altogether, these results shed light into a novel, yet poorly studied mechanism of cancer stemness induction by immunosuppressive myeloid cells involving active membrane-bound TGF-β1.

## Discussion

The role of suppressive myeloid cells in cancers has been extensively studied and characterized by a mounting body of literature over the last decade^56–59^. Many studies have more recently addressed the broad phenotypic and functional diversity and heterogeneity of myeloid cell subsets based on multi-omics approaches^14,60–62^. Likewise, multiple reports have identified a vast variety of mechanisms underlying tumor-induced generation, expansion and recruitment of these cells in different tissues and the modalities by which they suppress virtually all steps of anti-cancer immune response^57,60–63^. Along these lines, evidence has been provided that beyond their cardinal immunosuppressive functions, many suppressive myeloid cell populations can exert multifaceted tumor-promoting functions resulting in enhanced tumor proliferation, survival, invasion and metastasis^14^. However, the concept that immunosuppressive myeloid cells may also participate to cancer development by fostering the acquisition and maintenance of cancer cells with stemness properties has drawn much less scrutiny. Specifically, it has been proposed that TAM may support a stemness phenotype of cancer cells by activating signaling pathways such as STAT3, WNT/β-catenin or NF-κB, promoting CSC maintenance, EMT and therapy resistance^64–67^. Some reports have suggested that MDSC may also enhance the survival and self-renewal potencies of CSC through signalling pathways involving STAT3, NF-κB or CXCL2-CXCR2 and by increasing EMT markers^24,64,68–71^. However, the mechanistic bases underlying cancer-stemness induction by suppressive myeloid cells remained largely understudied and thus needed to be fully comprehended. In this study, we demonstrate that immunosuppressive cells generated from human circulating monocytes, HuMoSC^29^, and immunosuppressive myeloid cells isolated from breast cancer patients promote the acquisition of cancer stemness properties, characterized by different approaches, in tumor cell populations. The ability of cancer cells to form tumorspheres in selective medium condition, a functional hallmark of CSC, is enhanced by suppressive myeloid cells and coherently, the frequency of CSC in the breast cancer cell population is augmented after coculture with suppressive myeloid cells. Interestingly, only myeloid cells endowed with immunosuppressive properties, but not their non-immunosuppressive monocyte counterparts, were capable of promoting cancer stemness.

Different subsets of breast CSC, associated with epithelial (E-CSC) or mesenchymal (M-CSC) characteristics have been recently described^34,72–75^. One of the key finding of our current study is the peculiar capacity of suppressive myeloid cells to specifically promote the emergence of mesenchymal-like cancer stem cells expressing a dedicated gene expression pattern^34^. This is an important information as it has been reported that mesenchymal-like CSC, which are primarily localized at the edge of tumors, exhibit higher invasive and migratory properties, leading to more aggressive metastatic behavior. This plasticity of CSC dictated by the EMT program thus allows their differentiation through a broad spectrum of cell states (from fully epithelial to fully mesenchymal states) and governs their adaptability during tumor progression and metastasis^76^. The capability of suppressive myeloid cells to promote M-CSC occurrence may thus represent one additional mechanism by which these cells enhance breast cancer invasion and metastasis. However, whether HuMoSC polarize E-CSC into M-CSC through increased EMT activation or whether HuMoSC foster *de novo* M-CSC emergence from non-CSC remains to be determined. This is particularly relevant as the “mesenchymal subtype” of triple-negative breast cancers, characterized by the expression of EMT-related molecular markers, is a highly aggressive cancer with poor prognosis that correlates with metastatic dissemination and resistance to therapies^77^.

Suppressive myeloid cells are composed of multiple subsets characterized by more or less overlapping phenotypes and functions. Our data suggest that immunosuppressive myeloid cells expressing CD52 are primarily capable of promoting cancer stemness. The value of CD52 as an identifying marker for such population remains debatable as its expression has been detected on various immune cells (lymphocytes, monocytes, macrophages and dendritic cells)^78–81^. However, CD52 expression on T cells has been described as an immunoregulator in autoimmune diseases, either by release of its soluble form^82^, or by its membrane-bound form which may act as a costimulatory molecule for regulatory T cells^83^. Only few reports are available on the role of CD52 in myeloid cells, but recent studies have described CD52 as a marker of PMN-MDSC in cancer patients, further linking this molecule to an immunosuppressive function^84^. A recent report in cirrhosis patients described CD52-monocytes as endowed with immunosuppressive functions^85^. Conversely, CD52^+^ T and B cells have conversely been associated with better outcome in breast cancers^80,86^. The role of CD52^+^ myeloid cells in cancer promotion thus requires additional investigation. CD52 holds potential as a valuable marker for patient-derived monocytic, immunosuppressive, and stemness-inducing myeloid cells; however, its utility should be considered in conjunction with the expression of additional markers.

Mechanistically, we determined that breast cancer stemness induction by suppressive myeloid cells requires a direct cell-to-cell contact, thus the engagement of membrane-bound factors present at the surface of cancer cells and myeloid cells. Combined scRNAseq transcriptomic and an original cell surface proteomic approaches uncovered TGF-β1 as one potential molecule involved in the communications between suppressive myeloid cells and cancer cells. Blockade experiments further evidenced the importance of the TGFβ pathway for cancer stemness induction by suppressive myeloid cells. However, the culture supernatants of suppressive myeloid cells or myeloid cells and cancer cells co-cultures did not induce cancer cell stemness properties Furthermore, the levels of TGF-β1 were not significantly different in the culture supernatants of suppressive myeloid cells compared to that of monocytes that were not able to significantly foster cancer stemness. Conversely, TGF-β1 was detected in the membrane protein fraction of suppressive myeloid cells. Taken together, these observations point to a possible yet unknown role of membrane-bound TGF-β1 in breast cancer stemness induction. TGF-β1 is a multifunctional cytokine involved in a plethora of cellular processes and regulatory networks in physiological and pathological conditions^87–89^. It specifically plays an essential role in immune regulation^46^. TGF-β is initially synthesized as a precursor protein that is cleaved intracellularly to produce mature TGF-β and the latency-associated peptide (LAP), forming the small latent complex (SLC). This SLC is secreted and deposited in the extracellular matrix and activation of latent TGF-β involves release of the mature cytokine from LAP, which can occur through different mechanisms including proteolysis, pH changes, reactive oxygen species, and interactions with integrins or thrombospondin-1^46,90^. Importantly, TGFβ1 has been identified as one of the primary factors participating to cancer stemness induction^45,46,91^. However, the vast majority of the literature has evaluated the role of its soluble secreted form and only a few reports have documented the possibility that this molecule may function as a membrane-bound factor. Membrane-bound TGF-β has been detected on Tregs^48,50,51^, malignant B cells in B-cell non-Hodgkin lymphoma^49^ and colorectal cancer cells^53^ and was associated with immune regulation and cancer progression. To our knowledge, only one study has reported on the expression of membrane-bound TGF-β by murine suppressive myeloid cells, where it was shown to foster Treg polarization^47^.

Different hypotheses may explain the potential advantages of membrane-bound TGF-β over its soluble form. Membrane-bound TGF-β could circumvent extracellular matrix sequestration of the secreted latent complex TGF-β, impacting its activation and bioavailability^92,93^. This alternative mechanism may allow immunosuppressive myeloid cells to maintain a TGF-β signaling to cancer cells despite extracellular matrix regulatory effects. Similarly, this strategy may ensure that even if soluble TGF-β is depleted in the environment, membrane-bound TGF-β remains available. Additionally, membrane-bound TGF-β may differentially affect downstream signaling pathways compared to its soluble counterpart, promoting distinct phenotypic plasticity in CSC, such as enhancing mesenchymal traits or impacting their proliferative or migratory capacities.

Our discovery that membrane-bound TGF-β may act as an inducer of cancer stemness phenotype in breast cancer represents a potentially novel and clinically relevant finding. While secreted TGF-β is already known to foster stemness, its involvement in numerous physiological processes, both pathological and non-pathological, makes it a challenging therapeutic target. Current anti-TGF-β therapies predominantly target the secreted form, but targeting membrane-bound TGF-β may provide a more effective therapeutic approach more selectively focusing on cells responsible for cancer stemness promotion and maintenance. Additionally, the reported association of membrane-bound TGF-β with different immunosuppressive cells such as Tregs suggest that targeting this isoform may potentially reduce simultaneously immunosuppression and stemness induction, thereby achieving dual therapeutic benefits. However, the extent to which direct cell-to-cell contact and the membrane-bound TGF-β isoform are required in patient’s immunosuppressive myeloid cells fostering of CSC remains to be clarified. Finally, identifying the proteins associated with membrane-bound TGF-β on the cell membrane could reveal new therapeutic targets, potentially allowing for the development of new strategies to inhibit both the TGF-β pathway and its associated mechanisms more effectively.

## Acknowledgments

This project was supported by the French National League Against Cancer (National and Dordogne Committees), the SIRIC-BRIO (C.D., N.L.), the Foundation ARC (C.B.), The “Réseau Impulsion Newmoon”, and ITMO Aviesan grant (C.V. and N.L. (22CE070-00)); D.C., J.G. (22CE052-00). A.M., A.T-L. and T.B. are recipients of a MESRI doctoral funding. M.M. is recipient of a Cancer Biology Graduate Program (UB grad 2.0) doctoral funding (state support managed by the French National Research Agency (ANR) under reference ANR-20-SFRI-0001). D.A. received an Idex Visiting Scholar Award for Bordeaux University. C.B. is recipient of the GILEAD Research Scholars Programs in Oncology. We thank Anissa Zaafour, Anne-Claire Toublanc and Justine Vaché for technical assistance and Xavier Gauthereau from TBMCore OneCell facility for the help with single cell RNA sequencing sample preparation and technical advice. We also thank Scilicum company for BRB-sequencing and their help in the analysis, our collaborator Explicyte and especially Alban Bessede and all physicians of the Bergonié Institute in the identification and collection of patients samples.

## Author contributions

Conceptualization, T.B., C.B., C.V., C.D., N.L.; Formal analysis, T.B., C.B., D.C., C.D., N.L; Investigation, T.B., C.B., D.C., M.M., J.B., A.M., J.G., A.T-D., F.P., L.S., C.K., S.M., E.B.; Writing – original draft, T.B., C.B., C.D., N.L.; Supervision C.V., C.D., N.L.; Funding acquisition C.B., C.V., C.D., N.L.; Resources S.A., A.C., A.Z., G.M., P.R., B.L., B.B.; Data discussion, B.L., B.B., D.A.; All of the authors commented on the manuscript.

## Declaration of interests

The authors declare no competing interests.

## Material and Methods

### Lead contact

Further information and request for resources and reagents should be directed to and will be fulfilled by the lead contact, Nicolas Larmonier.

### Materials availability

This study did not generate new unique reagents.

### Data and code availability

Scripts used to process the data and generate the figures are available upon request. Any additional information required to reanalyze the data reported in this paper is available from the lead contact upon request.

## EXPERIMENTAL MODELS AND SUBJECT DETAILS

### Human specimens

A total of eight malignant pleural effusions collected between 2022 and 2024 from patients with invasive luminal or triple negative breast cancer at Bergonié Institute (Bordeaux, France) were included in this study, in accordance with institutional ethical guidelines and after informed consent of patient was obtained in accordance with the Declaration of Helsinki. The protocol was approved by the Ethics Committee of the Bergonié Institute. Seven out of eight pleural effusions contained sufficient CD45^+^ immune cells for FACS sorting and follow up experiments.

### Cell lines

MCF7 human Luminal A breast cancer cell line was purchased from the ATCC. ZsGreen-expressing MCF7 cells were generated using the ZsGreen lentiviral vector pRRLSIN-MND-LUC-IRES2-ZsGreen-WPRE purchased from VectUB (Vectorology platform, University of Bordeaux, France). MCF7 cells were cultured in DMEM/F12 plus GlutaMAX medium (Gibco) complemented with 10% heat-inactivated FBS (Gibco) and 1% Penicillin/Streptomycin (Gibco) at 37°C in 5% CO_2_.

### Lentiviral infection

1.10^5^ MCF7 cells at 30% confluency were infected with lentiviruses (at a Multiplicity of Infection of 20) for 24 hours in complete medium supplemented with 8 μg/mL Polybrene Infection / Transfection Reagent (Sigma-Aldrich). After 24 hours, lentiviruses were washed, and fresh medium was added. ZsGreen-expressing MCF7 cells (95% of total cells) were FACS sorted and expanded for further experiments.

### HuMoSC generation

HuMoSC were generated according to a published protocol^29^. Briefly, peripheral blood mononuclear cells (PBMC) were isolated from buffy coats of healthy blood donors using Ficoll density gradient centrifugation (**Figure S1A**). Monocytes were then isolated from PBMC by magnetic cell sorting using the AutoMACS^®^ Pro Separator and Human CD14^+^ cell isolation kit (Miltenyi Biotec) following manufacturer’s indications. Monocytes (1.10^6^ cells/mL) were incubated for 7 days in RPMI 1640 complemented with 10% FBS, 1% Glutamine, 1% sodium pyruvate, 1% Hepes, 1% non-essential amino acids, 1% Penicillin/Streptomycin (Gibco), recombinant human GM-CSF (10 ng/mL, Miltenyi Biotec) and IL-6 (10 ng/mL, Miltenyi Biotec) for 7 days. Sixty percent of the medium was replaced every 3 days with cytokines renewal. Semi-adherent HuMoSC were retrieved by Accutase cell detachment (Biolegend).

### Myeloid cell isolation from pleural effusion of breast cancer patients

Malignant pleural effusions were diluted 1:1 in PBS and filtered through a wide-mashed strainer followed by a 70μm cell strainer. Samples were then centrifuged at 500×g for 15 min, washed with PBS and centrifuged at 800×g for 15 min. Red Blood Cells were lysed in ACK for 4 min, washed with PBS and centrifuged 800×g for 10 min. Total cells were then counted and stained with CD33 (BC Biosciences) and CD52 (Miltenyi Biotec) antibodies and viability dye. After FSC/SSC doublets and dead cells removal, cells differentially expressing CD33 and CD52 were sorted (BD FACS Melody Cell Sorter, BD Biosciences).

## METHOD DETAILS

### Co-culture experiments

Classical 2D co-cultures of MCF7 and HuMoSC were conducted for 72 hours in 6 well plates. Cells were seeded at a 1:5 ratio (MCF7 : HuMoSC) at a density of 1.10^5^ cells/mL in MCF7 complete medium. When indicated, cells were treated with 5 μM TGF-βR type I/II inhibitor Ly2109761 (MedChemExpress) or TGF-β1,2,3 antibody (MAB1835, R&D Systems) at neutralizing concentration (1 μg/mL).

### Tumorsphere formation assay

MCF7 (300 cells per well) were seeded in flat-bottom 96-well plates, previously coated with 10% poly-2-hydroxymethyl methacrylate (polyHEMA, Sigma Aldrich) to prevent cell adhesion. Cells were incubated (37°C, 5% CO_2_) in serum free DMEM/F12 plus GlutaMAX complemented with 0.3% glucose, 1% N2-supplement (Thermo Fisher Scientific), 5 μg/mL insulin, 20 ng/mL human epithelial growth factor and 20 ng/mL basic fibroblast growth factor (Sigma-Aldrich). When indicated, monocytes, HuMoSC or patient’s myeloid cells were added in each well at a 1:5 ratio (1.500 cells per well) and wells were treated with 5 μM TGF-βR type I/II inhibitor Ly2109761 (MedChemExpress) or TGF-β1,2,3 antibody (MAB1835, R&D Systems) at neutralizing concentration (1 μg/mL). Ten wells per condition were seeded. The total number of tumorspheres were counted 3 or 6 days after seeding using an inverted light microscope (Nikon, Eclipse Ts2, objective ×20). Normalized tumorsphere forming efficiency was calculated as a ratio of the mean of all wells from the corresponding control condition.

### Live video-microscopy

The IncuCyte S3 Live-Cell Analysis System was used to analyze time-lapse tumorsphere formation of MCF7 cells in culture with HuMoSC. 300 MCF7 cells stably expressing ZsGreen Fluorescent protein were seeded per well and cultured for 7 days in tumorsphere forming conditions. HuMoSC or Monocytes (1.500 cells per well) were seeded after staining with CellTrace Far Red (Thermo Fisher Scientific), according to manufacturer’s indications. Throughout the assay, both phase and fluorescent images were collected using phase contrast, Green (300 ms exposure) and Red (400 ms exposure) fluorescence channels with a 10× lens and the S3/SX1 G/R Optical Module. 4 images per well were taken every 4 hours for 7 days, and each condition was run in ten replicates (10 wells per conditions). Images were analyzed, and quantitative data were generated using the Incucyte software (IncuCyte GUI Version 2021C). For final videos, the four images from the same well were fused in one single image using an in-house Python script through Jupiter Notebook v.6.5.2.

### Extreme Limiting Dilution Analysis (ELDA)

The tumorsphere ELDA was performed in similar conditions as previously published. Briefly, decreasing densities of MCF7 cells alone or after co-culture with HuMoSC were seeded in tumorsphere forming conditions (300, 100, 50, 30, 10, 5 cells per well) with the assumption that a single cancer stem-like cell can generate a tumorsphere. After determining the number of positive and negative wells (i.e. wells with at least one tumorsphere or no tumorsphere, respectively), we used the ELDA software^94^ for statistical analysis.

### Human T cell suppression assay

Myeloid cells immunosuppressive functions were determined by assessing their ability to suppress T-cell proliferation *in vitro*. Total T cells were purified from healthy blood donors PBMC by magnetic cell sorting using the AutoMACS^®^ Pro Separator and human Pan T Cell Isolation Kit (Miltenyi Biotec). The isolated lymphocytes were stained with CellTrace™ Violet (CTV, 5μM/106 cell/mL, 20 min at 37°C) (LifeTechnologies). Labeled T-cells were then activated or not with anti-CD3/CD28-coated beads (Dynabeads, Life Technologies) and plated in 96 well plates (8.0×10^4^ cells/well). Generated HuMoSC or patients’ myeloid cells were then added at the indicated ratios. T-cells proliferation was detected after 5 days by Flow Cytometry (Canto II cytometer, BD Biosciences) and analyzed using the proliferation module of the FlowJo (version 10.10.0) software.

### Transwell assay

MCF7 cells (1.10^5^ cells/well, 600 μL) were seeded in 24-well plates in complete medium and left in culture for two hours until adherent. A 0.3 μm pore size membrane transwell chamber (Corning) was then inserted and HuMoSC (5.10^5^ cells/insert, 100 μL) were added. Conditioned medium (CM) from HuMoSC or from previous 2D-co-culture of MCF7 and HuMoSC were used as controls.

### Live Immunofluorescence on tumorspheres

ZsGreen-expressing MCF7 (10.000 cells) and HuMoSC (50.000 cells) were co-cultured for 7 days in polyHEMA-coated non-adherent 6-well plates as described above. Cells were collected and incubated with APC-conjugated anti-CD14 and FITC-conjugated anti-EPCAM antibodies (STEMCELL Technologies) in an ice-cold buffer (PBS, 0.5% BSA, 2 mM EDTA) for 25 min at 4 °C. Cells were washed twice and incubated for 30 min at room temperature with Hoescht (Thermo Fisher Scientific) to stain nuclei. Live immunofluorescence acquisition was carried out with an Eclipse 50i epi-fluorescence microscope (Nikon) using the NIS-BR acquisition software (×40 objective, numerical aperture, 1.3). Images were analyzed using the NisElement Viewer software (Nikon Instruments Inc.).

### TGF-β1 quantitation

To quantitatively detect secreted TGF-β1, supernatant from HuMoSC generation at each day of their polarization (from Day 0 (monocyte) to Day 7 (HuMoSC)) as well as supernatant from MCF7 cells cultured alone and 72 hours 2D-co-culture of MCF7 and HuMoSC, both treated or not with Ly2109761 TGF-βRI/II inhibitor (10μM) were collected in 3 independent experiments. HuMoSC and MCF7 basal complete medium was also collected after the same incubation times as the cell to serve as reference quantity for TGF-β1 detection. Sandwich ELISA with the Human TGF-β1 DuoSet ELISA kit (R&D Systems) were performed according to manufacturer’s instructions. Optical Density was read at 450 nm with wavelength correction at 540 nm using a CLARIOstar PLUS microplate reader (BMG Labtech).

### Flow Cytometry staining

After MCF7 and HuMoSC co-culture as previously described, cells were stained with Aldefluor assay (STEMCELL Technologies) according to manufacturer’s guidelines along with CD45 (BD Biosciences) and live/Dead marker (Thermo Fisher Scientific) staining for discrimination of live cancer cells and HuMoSC. Additionally, another batch of cells were stained with CD45, CD24 and CD44 antibodies (BD Biosciences) and live/Dead marker (Thermo Fisher Scientific).

## BULK RNA BARCODING (BRB) SEQUENCING AND ANALYSIS

### RNA extraction and library preparation

MCF7 alone or MCF7 co-cultured for 24 with HuMoSC were sorted and subjected to RNA extraction for BRB sequencing. Total RNA was extracted using an RNeasy Minikit (Qiagen) following manufacturer’s instructions. RNA quantity was assessed using a NanoDropTM 8000 Spectrophotometer (Thermo Fisher Scientific), and RNA quality using a 2100 Bioanalyzer Instrument (Agilent Technologies, CA, USA). Only samples with an RNA integrity number (RIN) score >9 were subjected to sequencing.

### Bulk RNA barcoding (BRB) library preparation and sequencing

The 3’ Bulk RNA Barcoding and sequencing (BRB-seq)^95^ experiments were performed as described in^96^. Briefly, a first step of reverse transcription and template switching reactions were performed using 4 µL total RNA at 2.5 ng/µL. Then cDNA was purified and double strand cDNAs were generated by PCR. The sequencing libraries were built by tagmentation using 50 ng of double strand cDNA with the Illumina Nextera XT Kit (Illumina) following the manufacturer’s recommendations. The resulting library was sequenced on a NovaSeq sequencer following Illumina’s instructions by the IntegraGen Company (https://integragen.com/fr/). Image analysis and base calling were performed using RTA 2.7.7 and bcl2fastq 2.17.1.14. Adapter dimer reads were removed using DimerRemover (https://sourceforge.net/projects/dimerremover/).

### Differential gene expression analysis

The sequencing data were first demultiplexed: for each read pair, the 6-bp unique sample-specific barcode found in the first read was used to assign the read pair to its sample of origin if the six bases were of sufficient quality (Q ≥ 10) or discarded otherwise. Then, second reads were trimmed using Cutadapt 3.5 and aligned to the reference transcriptome using BWA 0.7.17. The number of transcripts molecules for a given gene in a sample was estimated by counting the number of unique 10-bp molecular identifier (UMI, located in the first read) corresponding to the set of second reads aligned to this gene in the reference transcriptome. Differential gene abundance was then performed using the DeSeq2 package^97^. Briefly, genes with a very low expression or not detected in enough samples were discarded to increase statistical power. Statistically significant DEGs were identified by using the Wald test p-value implemented into the DESeq2 package (p-value adjusted using the Benjamini & Hochberg (BH) False Discovery Rate approach, p ≤ 0.05) and a at least a 1-fold change relative to the control groups.

### Gene Set enrichment Analysis (GSEA) and signaling pathway enrichment

GSEA was performed using the MSigDBGSEA software^98^ (version 4.3.2, https://www.gsea-msigdb.org/gsea/msigdb). Hallmark EMT (version 2023.2)^99^ gene set database was used to infer EMT signature. EMT and stemness related signaling pathways enrichment analysis was performed using the *enrichKEGG* function from clusterProfiler package.

## SINGLE-CELL RNA SEQUENCING DATA PROCESSING AND ANALYSIS

### Single-cell FACS sorting

Three experimental conditions were sorted for single-cell RNA-sequencing using FACS Aria II (BD Biosciences). A first “control” condition consisting in MCF7 cells alone or HuMoSC alone sorted based on viability staining (Thermo Fisher Scientific). A second “tumorsphere” condition consisting in three-days old MCF7 tumorspheres dissociated with Accumax™ (Sigma-Aldrich) prior to live/Dead staining. HuMoSC cultured for 3 days alone in tumorsphere forming experimental conditions were also retrieved, stained with Live/Dead marker and sorted. Finally, a third “co-culture” condition consisted in a co-culture of MCF7 and HuMoSC in tumorsphere forming conditions for three days as previously described. Cells were retrieved, dissociated using Accumax™, stained with CD45 antibodies and live/Dead marker and sorted. Doublet cells were excluded from all sorting based on width against forward and side scatter. Sorted cells were centrifuged at 350×g at 4°C for 5 min and counted with trypan blue dye before being processed. All sequenced conditions were conducted with a viability requirement >80%.

### Single cell RNA sequencing Experiment

10.000 MCF7 cells and 10.000 HuMoSC per conditions were loaded into a chip to form Gel Bead-in-Emulsion in the Chromium Controller. Single-cell libraries were generated using the Chromium Next GEM Single Cell 3′ reagent Kits v3.1 (10X Genomics) according to manufacturer’s guidelines. cDNA was amplified by 11 PCR cycles and 11 cycles were also performed for library preparation (single index PCR). Libraries were pooled and sequenced on an Illumina NextSeq 2000 P3.

### Single-cell RNA-Seq data processing, quality control and cleaning

scRNA-seq data processing is summarized in **Figure S2A**. Each of the three 10X Chromium single-cell Gene Expression data were processed using CellRanger software 7.1.0, including demultiplexing, read alignment on human reference genome assembly GRCh38 (refdata-gex-GRCh38-202-A), barcoding and counting of unique UMI. Raw UMI count matrices were imported in R environment to perform deeper quality control steps to exclude low-quality cells. Cells containing less than 500 and more than 9.000 detected features and cells with more than 60.000 counts were discarded. Moreover, cells expressing more than 12% of mitochondrial genes or more than 17% of ribosomal genes were eliminated. A supplementary criterion was added to exclude stressed cells based on a stress response score^36^. The stress response score was calculated using *AddModuleScore* function from Seurat package (version 5.1.0) with the following list of genes: *FOSB*, *FOS*, *JUN*, *JUNB*, *JUND*, *ATF3*, *EGR1*, *HSPA1A*, *HSPA1B*, *HSP90AB1*, *HSPA8*, *HSPB1*, *IER3*, *IER2*, *BTG1*, *BTG2*, *DUSP1*.

### Cell doublet detection and removal

Three different doublet predictions were performed using *DoubletFinder* (version 2.0.3)^100^, *scDblFinder* (version 1.7.7)^101^ and *scds* (version 1.8.0)^102^. A consensus method was applied (i.e., a cell was considered as a multiplet and discarded if identified in at least two of the three methods). Doublets were estimated at around 1.7%, 3.6% and 2.9% in the Control, Tumorsphere and Co-culture experimental conditions, respectively.

### Normalization and Data Integration

The three pre-processed scRNA-seq data were normalized using Seurat’s *NormalizeData* with normalization.method = “LogNormalize” before integration. Functions *PrepSCTIntegration* with 3,000 features, *FindIntegrationAnchors* with dims = 30 and reduction = «rpca», and *IntegrateData* from Seurat were used to integrate all 3 samples.

### Unsupervised clustering, dimensionality reduction and Data Visualization

Unsupervised clustering of 15.902 cells (8.643 HuMoSC and 7.259 MCF7), dimensionality reduction and visualization were performed with Seurat (version 5.1.0). *RunPCA* function was used with 30 dimensions. The optimized number of dimensions used for *RunTSNE* and *RunUMAP* functions was automatically calculated with an in-house script. *FindNeighbors* and *Findclusters* (res=0.25) function were used to predict the eight clusters described in the main text.

### Differential gene expression analysis

Differential gene expression analysis was applied on integrated HuMoSC and MCF7 clusters independently. We used the Seurat functions *FindVariableFeatures* (selection.method = vst, nfeatures = 2000) and *FindAllMarkers* (min.pct = 0.25, logfc.threshold = 0.25) to identify top 20 discriminating genes between clusters.

### Identification of signature genes

The “Mesenchymal Cell Differentiation” gene signature was obtained from Harmonize 3.0 Database(https://maayanlab.cloud/Harmonizome/gene_set/mesenchymal+cell+differentiation/GO+Biological+Process+Annotations+2023)^37^. The “MDSC” signatures was extracted from PangloDB database (https://panglaodb.se). To validate these signatures, scores were calculated using *AddModuleScore* and visualization was performed with *FeaturePlot* functions from Seurat.

### Statistical cluster enrichment test

To identify the proportional difference in cell populations between “Tumorsphere conditions” and “Tumorsphere co-culture” experiments in our dataset, we used the *scProportionTest* package (https://github.com/rpolicastro/scProportionTest/releases/tag/v1.0.0) as previously published^103^.

### Transcriptomic cellular communication network prediction

The intercellular ligand-receptor interactions among all clusters of the single cell experiment were analyzed by CellChat (version 1.6.1) R package^42^. It applies a method of pattern recognition using non-negative matrix factorization to uncover both the overarching communication patterns and crucial signals within distinct cell groups. The standardized expression matrix was imported to create the CellChat object via the *createCellChat* function. The predicted ligand-receptor interactions were calculated using *ComputeCommunProb* (raw.use=FALSE), *filterCommunication* (min.cells = 10) and *computeCommunProbPathway* functions. To identify global outgoing communication patterns, we used the *IdentifyCommunicationPatterns* function with k=4 patterns and results were visualized using river plots. Interaction between all clusters depending on specific pathways of interest were represented using *netVisual_chord_cell* function.

## SURFACEOME PROTEOMIC ANALYSIS

### Cell surface protein biotinylation

Cell surface protein biotinylation was performed on monocytes, HuMoSC or MCF7 cells, as previously described^43^. A first set of experiments were performed on monocyte from three different blood donors and their corresponding polarized HuMoSC after seven days of cytokine exposure, as described above. A second set of experiments were performed on three distinct generations of HuMoSC from three different blood donors and three 15 cm petri dishes of MCF7 thawed independently and passaged once. The immunosuppressive and tumorsphere inducing properties of the HuMoSC were confirmed for each generation. First, cells were chilled on ice to prevent endocytosis or degradation of certain surface proteins. Cells were washed twice with ice-cold biotinylation buffer (PBS pH 7.4, 0.5 mM MgCl_2_, 1 mM CaCl_2_) and incubated for 1h at 4°C with 1 mg/mL Sulfo-NHS-LC-Biotin EZ-Link™ (Thermo Fisher Scientific) resuspended in biotinylation buffer. The biotinylation reaction was quenched with biotinylation buffer containing 0.1 M glycine for 10 min at 4°C. Cells were washed twice with ice-cold biotinylation buffer and surface-biotinylated cells were lysed in Surfaceome Lysis Buffer (SLB, biotinylation buffer supplemented with 1% Triton X-100, 150 mM NaCl, 10 mM Tris HCl, 5 mM iodoacetamide (Sigma-Aldrich), 1× protease inhibitor (Roche), 1 mM Na_3_VO_4_ and 1 mM Phenylmethylsulfonyl fluoride) for 30 min at 4°C and cell debris and nuclei were removed by centrifugation (16.000 × g, 10min at 4°C). Total proteins were quantified using Pierce BCA Protein Assay Kit (Thermo Fisher Scientific) according to the manufacturer’s guidelines. For HuMoSC and monocytes, the three independent donors were pooled into one single aliquot and equally split in three replicates in order to prevent patient’s variation biases in the qualitative proteomic analysis. Proteins were stored in SLB at -80°C prior to streptavidin pull-down.

### Streptavidin pull-down of biotinylated proteins and peptide digestion

Biotinylated proteins were isolated from 7 mg (monocyte and HuMoSC surfaceome) or 10 mg (HuMoSC and MCF7 surfaceome) total protein by incubating cell lysates with high-capacity streptavidin agarose resin (Thermo Fisher Scientific) for 2h at 4 °C. Beads were washed extensively with intermittent centrifugation at 1000 × g for 5 min to eliminate all potential contaminants bound to biotinylated proteins. Three washes were performed with SLB, once with PBS pH 7.4/0.5% (w/v) sodium dodecyl sulfate (SDS), and then beads were incubated with PBS/0.5% SDS/100 mM dithiothreitol (DTT), for 20 min at RT. Further washes were performed with 6 M urea in 100 mM Tris–HCl pH 8.5, followed by incubation with 6 M urea/100 mM Tris–HCl pH 8.5/50 mM iodoacetamide, for 20 min at RT. Additional washes were performed with 6 M urea/100 mM Tris– HCl pH 8.5, PBS pH 7.4 and then water. For proteomic analysis, beads were rinsed thrice with 50 mM ammonium bicarbonate (NH_4_HCO_3_) pH 8.5, and re-suspended in 400 μL of 50 mM NH_4_HCO_3_ pH 8.5 containing 4 μg of proteomics grade trypsin (Sigma-Aldrich), overnight at 37 °C. The proteins were further digested with an additional 4 μg trypsin for 4 h at 37 °C. The resulting tryptic peptides were then collected by centrifugation at 10,000 × g, for 10 min at RT. The beads were washed twice with MS grade water and the tryptic fractions pooled. The tryptic fractions were dried to completion in a SpeedVac and re-suspended in MS solvent (5% aqueous Acetonitrile (ACN), 0.2% FA).

### Liquid Chromatography double tandem mass-spectrometry (LC-MS/MS)

Samples were loaded on a 1.5 µL C18 pre-column (Optimize Technologies) connected directly to the switching valve. Samples were separated on a homemade reversed-phase column (150 μm i.d. by 150 mm) with a 56-min gradient from 10 to 30% ACN/0.2% FA and a 600-nl/min flow rate on a Ultimate 3000 LC system (Eksigent) connected to a Q-Exactive Plus (Thermo Fisher Scientific). Each full MS spectrum acquired at a resolution of 70.000 was followed by 12 tandem-MS (MS– MS) spectra on the most abundant multiply charged precursor ions. Tandem-MS experiments were performed using collision-induced dissociation (CID) at a collision energy of 27%. Proteomic samples were analyzed as biological, back-to-back triplicates per condition. Peptides were identified using PEAKS (v7.0, Bioinformatics Solutions) and peptide sequences were blasted against the corresponding Uniprot database. Mass tolerances on precursor and fragment ions were 10 ppm and 0.01 Da, respectively. The false discovery rate (FDR) for peptide and protein was set to 0.5%. The minimum number of peptides per protein was set to 2, and minimum peptide length was set to ∼6 amino acids. Search criteria included a static modification of cysteine residues of +57.0214 Da; a variable modification of +15.9949 Da to include potential oxidation of methionines; and a modification of +79.966 on serine, threonine, or tyrosine for the identification of phosphorylation.

### Surface Protein Identification and Data Filtering

Peptide precursor intensities were extracted using an integral algorithm of PEAKS software (version 8.05). Proteins were then selected based on their detection in all replicates of the same sample, with a minimum of two unique peptides, which were verified for uniqueness and leucine/isoleucine switch using the neXtProt checker^104^. These proteins were considered as identified with high confidence. Gene Ontology Cellular Component (GO.CC) terms and functional annotations were queried for all identified proteins using g:profiler (http://biit.cs.ut.ee/gprofiler/index.cgi) to get an objective estimation about the number of proteins specific to the cell surface. We then associated a SPAT (Surface Protein Annotation Tool) score^44^ for each detected protein using the public web portal (https://spat.leucegene.ca/, version 3.4). This score ranges from 0 to 10 and represents the probability of a protein to be located at the cell surface based on structural motifs and public proteomic databases. We kept for the analysis only proteins with a SPAT score ≥8, considered as high-confidence score for surface protein identification^105^.

### Interaction Network Construction and Differential Expression Analysis

To interrogate the potential surface protein candidates in the physical interaction between HuMoSC and MCF7, we extracted the filtered list of protein independently of their intensity and annotated them based on their origin (HuMoSC or MCF7). We used Talkien^106^ web tool (version v1.2, https://shiny.odap-ico.org/talkien/) to create a surface proteomic-based interaction network. For consistency and comparison with the transcriptomic prediction described above, we set the used Ligand-Receptor DataBase on CellChat only. For quantitative surfaceome comparison between monocytes and HuMoSC, proteins were considered differentially upregulated by HuMoSC if log2 FC (HuMoSC/Monocyte) values were ≥ 2, and downregulated if ≤ −2. Some proteins were only identified in HuMoSC (n=72) or Monocyte (n=56), making it impossible to calculate fold-change values. Therefore, we arbitrarily assigned +5 if proteins were only present in HuMoSC, and −5 if they were only identified in monocytes. These values were chosen as it was higher than our highest and lowest HuMoSC/Monocyte fold-change, respectively.

## STATISTICAL ANALYSIS

Tests used for statistical analyses are described in the figure legends and have been determined using GraphPad Prism v8. Symbols for significance: ns, non-significant; ∗, <0.05, ∗∗, <0.01; ∗∗∗, <0.001; ∗∗∗∗, <0.0001. Values were expressed as mean ± S.E.M.

## SUPPLEMENTAL INFORMATION TITLES AND LEGENDS

**Figure S1. Immunosuppressive myeloid cells foster cancer stemness. Related to Figure 1**

(A) Workflow of in vitro HuMoSC generation from healthy blood donor monocytes. Kinetics of immunosuppression and tumorsphere formation were performed each day as depicted.

(B) Representative images of the obtained MCF7 tumorspheres alone (left) or in co-culture with HuMoSC (right)

(C) Flow cytometry gating strategy of CD33-expressing myeloid cells from pleural effusion of breast cancer patients.

(D-E) Normalized T cell proliferation (D) and MCF7 tumorsphere formation assay (E) after co-culture with HuMoSC at each day of their polarization from monocytes (+D0) to HuMoSC (+D7).

**Figure S2. Cluster annotation from MCF7 and HuMoSC scRNAseq integrated UMAP. Related to Figure 2**

(A) Pipeline of ingle cell RNA-seq bioinformatic analyses

(B) Feature Plot of *PTPRC* expression for immune cell and cancer cell identification.

(C) UMAP visualization colored according to cell cycle Phases determined with the Cell Cycle package.

(D) Feature Plot of MKI67 and STMN1 expression as assessment of proliferating cells.

**Figure S3. CD52 expression by myeloid cells. Related to Figure 3**

(A) Flow cytometry analysis of CD52 expression on HuMoSC (blue, CD52^+^; green, CD52^-^) and monocytes (grey).

(B) Flow cytometry analysis of CD33 and CD52 expressing cells isolated from pleural effusion of breast cancer patients.

**Figure S4. Predictive identification of putative interacting molecular partners between myeloid cells and cancer cells. Related to Figure 4.**

(A) Tumorsphere formation live videomiscroscopy quantification of ZsGreen and CellTrace Red overlay intensity over time, as assessment of cellular proximity, of MCF7 cells alone (Control) or in co-culture with HuMoSC or monocyte. Dashed line: 24h timepoint.

(B) Kernel density plot showing log2 intensity of ACVR1 in MCF7 surfaceome. EPCAM was used as reference for protein intensities. Dashed line: calculated standard deviation (SD) of the three replicates.

**Figure S5. Membrane-bound TGF-β is required for cancer stemness induction. Related to Figure 5.**

Tumorsphere formation assay of MCF7 cells alone treated or not with 2.5 ng/mL TGF-β1, 5 μM TGF-βI/II inhibitor Ly2109761, 1 μg/mL αTGF-β neutralizing antibody or the combination of TGF-β1 and αTGF-β. Data are means ± S.E.M. ∗∗∗∗p < 0.0001. NS, not significant. Kruskal-Wallis test with Dunn’s multiple comparisons post-test. Data are means of n = 3 independent biological replicates.

**Supplemental Table 1. HuMoSC and MCF7 surfaceome**

**Supplemental Table 2. HuMoSC and Monocyte surfaceome**

**Supplemental Table 3. Comparative table of HuMoSC and monocyte surfaceome**

**Supplemental video 1. Time lapse videomicroscopy (Incucyte) analysis of MCF tumorsphere formation**

**Supplemental video 2. Time lapse videomicroscopy (Incucyte) analysis of MCF tumorsphere formation in presence of HuMoSC**

**Supplemental video 3. Time lapse videomicroscopy (Incucyte) analysis of MCF tumorsphere formation in presence of Monocytes**

## Notes

### Competing Interest Statement

The authors have declared no competing interest.

## References

1. Hanahan, D. Hallmarks of Cancer: New Dimensions. Cancer Discov 12, 31–46 (2022).

2. Gabrilovich, D. I., Ostrand-Rosenberg, S. & Bronte, V. Coordinated regulation of myeloid cells by tumours. Nature Reviews Immunology *2012 12:4* 12, 253–268 (2012).

3. Pan, Y., Yu, Y., Wang, X. & Zhang, T. Tumor-Associated Macrophages in Tumor Immunity. Front Immunol 11, 583084 (2020).

4. Gabrilovich, D. I. & Nagaraj, S. Myeloid-derived suppressor cells as regulators of the immune system. Nature Reviews Immunology *2009 9:3* 9, 162–174 (2009).

5. Davidov, V., Jensen, G., Mai, S., Chen, S. H. & Pan, P. Y. Analyzing One Cell at a TIME: Analysis of Myeloid Cell Contributions in the Tumor Immune Microenvironment. Front Immunol 11, (2020).

6. Vonderheide, R. H., Domchek, S. M. & Clark, A. S. Immunotherapy for Breast Cancer: What Are We Missing? Clin Cancer Res 23, 2640–2646 (2017).

7. Gatti-Mays, M. E. et al. If we build it they will come: targeting the immune response to breast cancer. NPJ Breast Cancer 5, 37 (2019).

8. Harbeck, N. et al. Breast cancer. Nat Rev Dis Primers 5, (2019).

9. Alizadeh, D. & Larmonier, N. Chemotherapeutic targeting of cancer-induced immunosuppressive cells. Cancer Res 74, 2663 (2014).

10. Alizadeh, D. et al. Doxorubicin eliminates myeloid-derived suppressor cells and enhances the efficacy of adoptive T-cell transfer in breast cancer. Cancer Res 74, 104–118 (2014).

11. Zhang, W. jie et al. Tumor-associated macrophages correlate with phenomenon of epithelial-mesenchymal transition and contribute to poor prognosis in triple-negative breast cancer patients. Journal of Surgical Research 222, 93–101 (2018).

12. Law, A. M. K., Valdes-Mora, F. & Gallego-Ortega, D. Myeloid-Derived Suppressor Cells as a Therapeutic Target for Cancer. Cells 9, (2020).

13. Akkari, L. et al. Defining myeloid-derived suppressor cells. Nat Rev Immunol (2024) doi:10.1038/S41577-024-01062-0.

14. Blaye, C., Boyer, T., Peyraud, F., Domblides, C. & Larmonier, N. Beyond Immunosuppression: The Multifaceted Functions of Tumor-Promoting Myeloid Cells in Breast Cancers. Front Immunol 13, 838040 (2022).

15. Qian, B. Z. & Pollard, J. W. Macrophage diversity enhances tumor progression and metastasis. Cell 141, 39–51 (2010).

16. Chen, P., Hsu, W. H., Han, J., Xia, Y. & DePinho, R. A. Cancer Stemness Meets Immunity: From Mechanism to Therapy. Cell Rep 34, (2021).

17. Zhang, B. et al. Macrophage-expressed CD51 promotes cancer stem cell properties via the TGF-β1/smad2/3 axis in pancreatic cancer. Cancer Lett 459, 204–215 (2019).

18. Hide, T. et al. Oligodendrocyte Progenitor Cells and Macrophages/Microglia Produce Glioma Stem Cell Niches at the Tumor Border. EBioMedicine 30, 94–104 (2018).

19. Wan, S. et al. Tumor-associated macrophages produce interleukin 6 and signal via STAT3 to promote expansion of human hepatocellular carcinoma stem cells. Gastroenterology 147, 1393– 1404 (2014).

20. Nomura, A. et al. NFκB-Mediated Invasiveness in CD133+ Pancreatic TICs Is Regulated by Autocrine and Paracrine Activation of IL1 Signaling. Mol Cancer Res 16, 162–172 (2018).

21. Guo, L. et al. Induction of breast cancer stem cells by M1 macrophages through Lin-28B-let-7-HMGA2 axis. Cancer Lett 452, 213–225 (2019).

22. Lu, H. et al. A breast cancer stem cell niche supported by juxtacrine signalling from monocytes and macrophages. Nat Cell Biol 16, 1105–1117 (2014).

23. Raghavan, S., Mehta, P., Xie, Y., Lei, Y. L. & Mehta, G. Ovarian cancer stem cells and macrophages reciprocally interact through the WNT pathway to promote pro-tumoral and malignant phenotypes in 3D engineered microenvironments. J Immunother Cancer 7, (2019).

24. Ouzounova, M. et al. Monocytic and granulocytic myeloid derived suppressor cells differentially regulate spatiotemporal tumour plasticity during metastatic cascade. Nat Commun 8, (2017).

25. Ai, L. et al. Myeloid-derived suppressor cells endow stem-like qualities to multiple myeloma cells by inducing piRNA-823 expression and DNMT3B activation. Mol Cancer 18, (2019).

26. Li, X. et al. Myeloid-derived suppressor cells promote epithelial ovarian cancer cell stemness by inducing the CSF2/p-STAT3 signalling pathway. FEBS J 287, 5218–5235 (2020).

27. Chen, P., Hsu, W. H., Han, J., Xia, Y. & DePinho, R. A. Cancer Stemness Meets Immunity: From Mechanism to Therapy. Cell Rep 34, (2021).

28. Reya, T., Morrison, S. J., Clarke, M. F. & Weissman, I. L. Stem cells, cancer, and cancer stem cells. Nature 414, 105–111 (2001).

29. Janikashvili, N. et al. Human monocyte-derived suppressor cells control graft-versus-host disease by inducing regulatory forkhead box protein 3–positive CD8+ T lymphocytes. Journal of Allergy and Clinical Immunology 135, 1614–1624.e4 (2015).

30. Lamouille, S., Xu, J. & Derynck, R. Molecular mechanisms of epithelial–mesenchymal transition. Nature Reviews Molecular Cell Biology *2014 15:3* 15, 178–196 (2014).

31. Clarke, M. F. et al. Cancer stem cells--perspectives on current status and future directions: AACR Workshop on cancer stem cells. Cancer Res 66, 9339–9344 (2006).

32. Garnache-Ottou, F. et al. Expression of the myeloid-associated marker CD33 is not an exclusive factor for leukemic plasmacytoid dendritic cells. Blood 105, 1256–1264 (2005).

33. Toor, S. M. et al. Differential gene expression of tumor-infiltrating CD33+ myeloid cells in advanced-versus early-stage colorectal cancer. Cancer Immunol Immunother 70, 803 (2021).

34. Liu, S. et al. Breast Cancer Stem Cells Transition between Epithelial and Mesenchymal States Reflective of their Normal Counterparts. Stem Cell Reports 2, 78 (2014).

35. Pastushenko, I. & Blanpain, C. EMT Transition States during Tumor Progression and Metastasis. Trends Cell Biol 29, 212–226 (2019).

36. Denisenko, E. et al. Systematic assessment of tissue dissociation and storage biases in single-cell and single-nucleus RNA-seq workflows. Genome Biol 21, (2020).

37. Rouillard, A. D. et al. The harmonizome: a collection of processed datasets gathered to serve and mine knowledge about genes and proteins. Database (Oxford) 2016, (2016).

38. Gulati, G. S. et al. Single-cell transcriptional diversity is a hallmark of developmental potential. Science (1979) 367, 405–411 (2020).

39. Bronte, V. et al. Recommendations for myeloid-derived suppressor cell nomenclature and characterization standards. Nature Communications *2016 7:1* 7, 1–10 (2016).

40. Rao, S. P. et al. Human Peripheral Blood Mononuclear Cells Exhibit Heterogeneous CD52 Expression Levels and Show Differential Sensitivity to Alemtuzumab Mediated Cytolysis. PLoS One 7, e39416 (2012).

41. Ratzinger, G., Reagan, J. L., Heller, G., Busam, K. J. & Young, J. W. Differential CD52 expression by distinct myeloid dendritic cell subsets: implications for alemtuzumab activity at the level of antigen presentation in allogeneic graft-host interactions in transplantation. Blood 101, 1422– 1429 (2003).

42. Jin, S. et al. Inference and analysis of cell-cell communication using CellChat. Nature Communications *2021 12:1* 12, 1–20 (2021).

43. Aubert, L. et al. Copper bioavailability is a KRAS-specific vulnerability in colorectal cancer. Nature Communications *2020 11:1* 11, 1–15 (2020).

44. Spinella, J. et al. SPAT: Surface Protein Annotation Tool. bioRxiv 2023.07.07.547075 (2023) doi:10.1101/2023.07.07.547075.

45. Oshimori, N. & Fuchs, E. The Harmonies Played by TGF-β in Stem Cell Biology. Cell Stem Cell 11, 751 (2012).

46. Batlle, E. & Massagué, J. Transforming Growth Factor-β Signaling in Immunity and Cancer. Immunity 50, 924–940 (2019).

47. Huang, M. T. et al. Short-chain fatty acids ameliorate allergic airway inflammation via sequential induction of PMN-MDSCs and Treg cells. Journal of Allergy and Clinical Immunology: Global 2, 100163 (2023).

48. Ostroukhova, M. et al. Tolerance induced by inhaled antigen involves CD4+ T cells expressing membrane-bound TGF-β and FOXP3. Journal of Clinical Investigation 114, 28 (2004).

49. Yang, Z. Z. et al. Soluble and Membrane-Bound TGF-β-Mediated Regulation of Intratumoral T Cell Differentiation and Function in B-Cell Non-Hodgkin Lymphoma. PLoS One 8, e59456 (2013).

50. Nakamura, K., Kitani, A. & Strober, W. Cell contact-dependent immunosuppression by CD4(+)CD25(+) regulatory T cells is mediated by cell surface-bound transforming growth factor beta. J Exp Med 194, 629–644 (2001).

51. Ghiringhelli, F. et al. CD4+CD25+ regulatory T cells inhibit natural killer cell functions in a transforming growth factor-beta-dependent manner. J Exp Med 202, 1075–1085 (2005).

52. Ahn, Y.-O., Lee, J.-C., Sung, M.-W. & Heo, D. S. Presence of membrane-bound TGF-beta1 and its regulation by IL-2-activated immune cell-derived IFN-gamma in head and neck squamous cell carcinoma cell lines. J Immunol 182, 6114–6120 (2009).

53. Baker, K., Raut, P. & Jass, J. R. Colorectal cancer cells express functional cell surface-bound TGFβ. Int J Cancer 122, 1695–1700 (2008).

54. He, X. et al. Mechanism of action and efficacy of LY2109761, a TGF-β receptor inhibitor, targeting tumor microenvironment in liver cancer after TACE. Oncotarget 9, 1130 (2018).

55. Fan, Q. M. et al. Tumor-associated macrophages promote cancer stem cell-like properties via transforming growth factor-beta1-induced epithelial-mesenchymal transition in hepatocellular carcinoma. Cancer Lett 352, 160–168 (2014).

56. Broz, M. L. & Krummel, M. F. The emerging understanding of myeloid cells as partners and targets in tumor rejection. Cancer Immunol Res 3, 313–319 (2015).

57. Gabrilovich, D. I., Ostrand-Rosenberg, S. & Bronte, V. Coordinated regulation of myeloid cells by tumours. Nature Reviews Immunology *2012 12:4* 12, 253–268 (2012).

58. Merad, M., Sathe, P., Helft, J., Miller, J. & Mortha, A. The Dendritic Cell Lineage: Ontogeny and Function of Dendritic Cells and Their Subsets in the Steady State and the Inflamed Setting. Annu Rev Immunol 31, 563–604 (2013).

59. Noy, R. & Pollard, J. W. Tumor-associated macrophages: from mechanisms to therapy. Immunity 41, 49–61 (2014).

60. van Vlerken-Ysla, L., Tyurina, Y. Y., Kagan, V. E. & Gabrilovich, D. I. Functional states of myeloid cells in cancer. Cancer Cell 41, 490–504 (2023).

61. Elliott, L. A., Doherty, G. A., Sheahan, K. & Ryan, E. J. Human Tumor-Infiltrating Myeloid Cells: Phenotypic and Functional Diversity. Front Immunol 8, (2017).

62. Dou, A. & Fang, J. Heterogeneous Myeloid Cells in Tumors. Cancers (Basel) 13, (2021).

63. Yang, Y., Li, C., Liu, T., Dai, X. & Bazhin, A. V. Myeloid-Derived Suppressor Cells in Tumors: From Mechanisms to Antigen Specificity and Microenvironmental Regulation. Front Immunol 11, 540749 (2020).

64. Chen, P., Hsu, W. H., Han, J., Xia, Y. & DePinho, R. A. Cancer Stemness Meets Immunity: From Mechanism to Therapy. Cell Rep 34, 108597 (2021).

65. Lu, H. et al. A breast cancer stem cell niche supported by juxtacrine signalling from monocytes and macrophages. Nat Cell Biol 16, 1105–1117 (2014).

66. Huang, R. et al. CCL5 derived from tumor-associated macrophages promotes prostate cancer stem cells and metastasis via activating β-catenin/STAT3 signaling. Cell Death Dis 11, (2020).

67. Sarkar, S. et al. Therapeutic activation of macrophages and microglia to suppress brain tumor-initiating cells. Nat Neurosci 17, 46–55 (2014).

68. Wang, Y. et al. Granulocytic Myeloid-Derived Suppressor Cells Promote the Stemness of Colorectal Cancer Cells through Exosomal S100A9. Adv Sci (Weinh) 6, (2019).

69. Kuroda, H. et al. Prostaglandin E2 produced by myeloid-derived suppressive cells induces cancer stem cells in uterine cervical cancer. Oncotarget 9, 36317–36330 (2018).

70. Panni, R. Z. et al. Tumor-induced STAT3 activation in monocytic myeloid-derived suppressor cells enhances stemness and mesenchymal properties in human pancreatic cancer. Cancer Immunol Immunother 63, 513–528 (2014).

71. Peng, D. et al. Myeloid-derived suppressor cells endow stem-like qualities to breast cancer cells through IL-6/STAT3 and NO/NOTCH cross-talk signaling. Cancer Res 76, 3156 (2016).

72. Lambert, A. W. & Weinberg, R. A. Linking EMT programmes to normal and neoplastic epithelial stem cells. Nature Reviews Cancer *2021 21:5* 21, 325–338 (2021).

73. Pastushenko, I. et al. Identification of the tumour transition states occurring during EMT. Nature *2018 556:7702* 556, 463–468 (2018).

74. Pastushenko, I. et al. Fat1 deletion promotes hybrid EMT state, tumour stemness and metastasis. Nature *2020 589:*7842 589, 448–455 (2020).

75. Grosse-Wilde, A. et al. Stemness of the hybrid Epithelial/Mesenchymal State in Breast Cancer and Its Association with Poor Survival. PLoS One 10, (2015).

76. Dongre, A. & Weinberg, R. A. New insights into the mechanisms of epithelial–mesenchymal transition and implications for cancer. Nature Reviews Molecular Cell Biology *2018 20:2* 20, 69–84 (2018).

77. Krakhmal, N. V., Babyshkina, N. N. & Vtorushin, S. V. Mesenchymal Subtype of Triple-Negative Breast Cancer: A Review of Specific Features. International Journal of Biomedicine 13, 14–19 (2023).

78. Kasarello, K. & Mirowska-Guzel, D. Anti-CD52 Therapy for Multiple Sclerosis: An Update in the COVID Era. Immunotargets Ther 10, 237–246 (2021).

79. Ginaldi, L. et al. Levels of expression of CD52 in normal and leukemic B and T cells: Correlation with in vivo therapeutic responses to Campath-1H. Leuk Res 22, 185–191 (1998).

80. Wang, J. et al. CD52 Is a Prognostic Biomarker and Associated With Tumor Microenvironment in Breast Cancer. Front Genet 11, (2020).

81. Shathili, A. M. et al. Specific sialoforms required for the immune suppressive activity of human soluble CD52. Front Immunol 10, 442865 (2019).

82. Bandala-Sanchez, E., Bediaga, N. G., Naselli, G., Neale, A. M. & Harrison, L. C. Siglec-10 expression is up-regulated in activated human CD4+ T cells. Hum Immunol 81, 101–104 (2020).

83. Bandala-Sanchez, E. et al. T cell regulation mediated by interaction of soluble CD52 with the inhibitory receptor Siglec-10. Nat Immunol 14, 741–748 (2013).

84. Pettinella, F. et al. Surface CD52, CD84, and PTGER2 mark mature PMN-MDSCs from cancer patients and G-CSF-treated donors. Cell Rep Med 5, (2024).

85. Geng, A. et al. Distinct circulating monocytes up-regulate CD52 and sustain innate immune function in patients with cirrhosis unless acute decompensation emerges. bioRxiv 2024.04.03.587894 (2024) doi:10.1101/2024.04.03.587894.

86. Ma, Y. F. et al. The immune-related gene CD52 is a favorable biomarker for breast cancer prognosis. Gland Surg 10, 78098–78798 (2021).

87. Massagué, J. TGFβ signalling in context. Nature Reviews Molecular Cell Biology *2012 13:10* 13, 616–630 (2012).

88. Zhao, B. & Chen, Y.-G. Regulation of TGF-β Signal Transduction. Scientifica (Cairo) 2014, 1–9 (2014).

89. Massagué, J. TGF-beta signal transduction. Annu Rev Biochem 67, 753–791 (1998).

90. Tzavlaki, K. & Moustakas, A. TGF-β Signaling. Biomolecules 10, (2020).

91. Heldin, C. H. & Moustakas, A. Signaling Receptors for TGF-β Family Members. Cold Spring Harb Perspect Biol 8, (2016).

92. Horiguchi, M., Ota, M. & Rifkin, D. B. Matrix control of transforming growth factor-β function. The Journal of Biochemistry 152, 321–329 (2012).

93. Chakravarthy, A., Khan, L., Bensler, N. P., Bose, P. & De Carvalho, D. D. TGF-β-associated extracellular matrix genes link cancer-associated fibroblasts to immune evasion and immunotherapy failure. Nature Communications *2018 9:1* 9, 1–10 (2018).

94. Hu, Y. & Smyth, G. K. ELDA: Extreme limiting dilution analysis for comparing depleted and enriched populations in stem cell and other assays. J Immunol Methods 347, 70–78 (2009).

95. Alpern, D. et al. BRB-seq: Ultra-affordable high-throughput transcriptomics enabled by bulk RNA barcoding and sequencing. Genome Biol 20, 1–15 (2019).

96. Giacosa, S. et al. Cooperative Blockade of CK2 and ATM Kinases Drives Apoptosis in VHL-Deficient Renal Carcinoma Cells through ROS Overproduction. Cancers (Basel) 13, 576 (2021).

97. Love, M. I., Huber, W. & Anders, S. Moderated estimation of fold change and dispersion for RNA-seq data with DESeq2. Genome Biol 15, (2014).

98. Subramanian, A. et al. Gene set enrichment analysis: A knowledge-based approach for interpreting genome-wide expression profiles. Proc Natl Acad Sci U S A 102, 15545–15550 (2005).

99. Liberzon, A. et al. The Molecular Signatures Database (MSigDB) hallmark gene set collection. Cell Syst 1, 417–425 (2015).

100. McGinnis, C. S., Murrow, L. M. & Gartner, Z. J. DoubletFinder: Doublet Detection in Single-Cell RNA Sequencing Data Using Artificial Nearest Neighbors. Cell Syst 8, 329–337.e4 (2019).

101. Germain, P. L., Robinson, M. D., Lun, A., Garcia Meixide, C. & Macnair, W. Doublet identification in single-cell sequencing data using scDblFinder. F1000Res 10, (2022).

102. Bais, A. S. & Kostka, D. scds: computational annotation of doublets in single-cell RNA sequencing data. Bioinformatics 36, 1150–1158 (2020).

103. Miller, S. A. et al. LSD1 and aberrant DNA methylation mediate persistence of enteroendocrine progenitors that support BRAF mutant colorectal cancer. Cancer Res 81, 3791 (2021).

104. Schaeffer, M. et al. The neXtProt peptide uniqueness checker: a tool for the proteomics community. Bioinformatics 33, 3471–3472 (2017).

105. Bordeleau, M. E. et al. Immunotherapeutic targeting of surfaceome heterogeneity in AML. Cell Rep 43, (2024).

106. Moratalla-Navarro, F., Moreno, V. & Sanz-Pamplona, R. TALKIEN: crossTALK IntEraction Network. A web-based tool for deciphering molecular communication through ligand-receptor interactions. Mol Omics 19, 688–696 (2023).

107. Hao, Y. et al. Dictionary learning for integrative, multimodal and scalable single-cell analysis. Nature Biotechnology *2023 42:2* 42, 293–304 (2023).

