## Supplementary figures and images for "Immunosuppressive myeloid cells induce mesenchymal-like breast cancer stem cells by a mechanism involving membrane-bound TGF-β1"

### Supplemental Figures 1-5

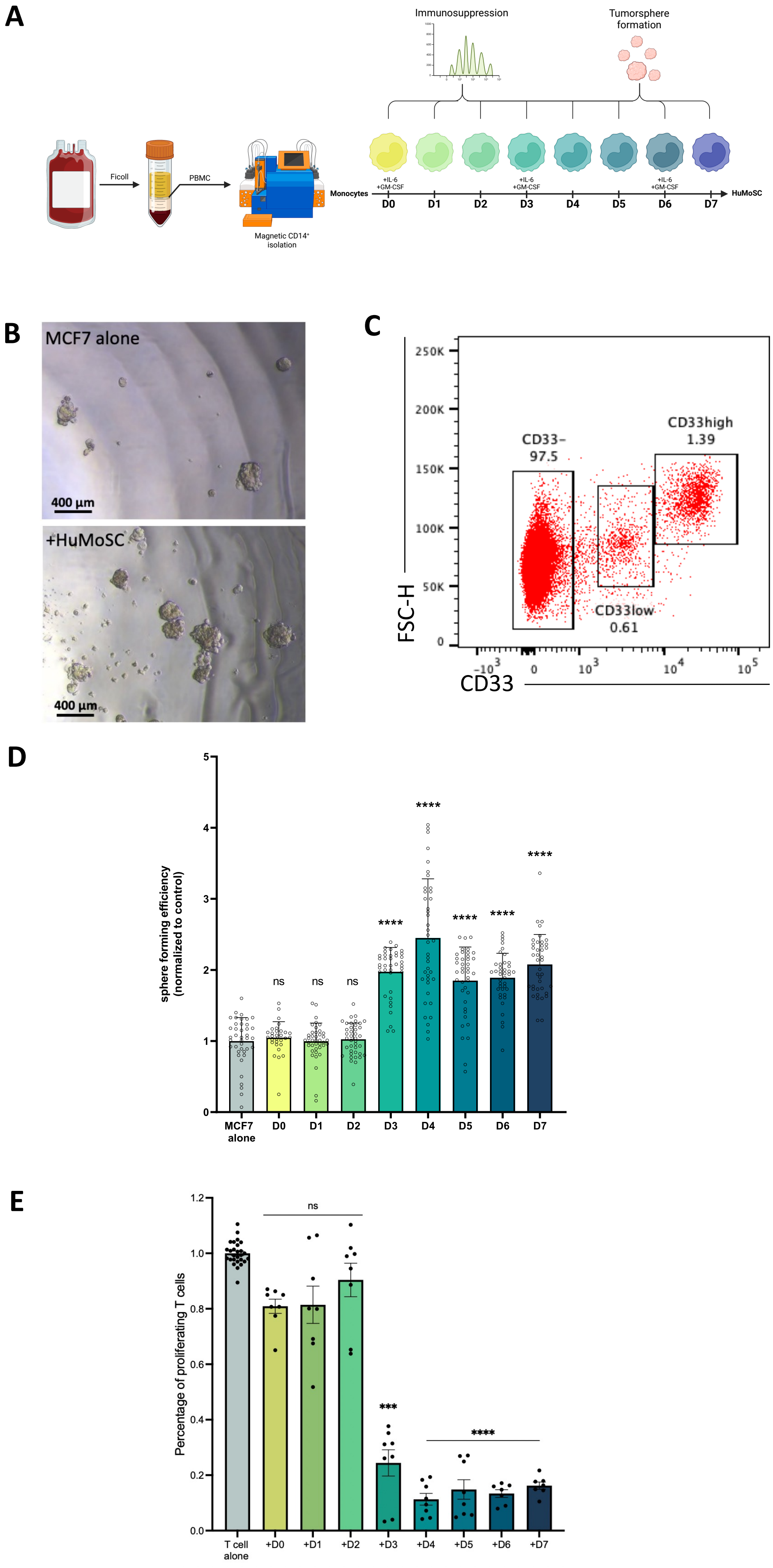

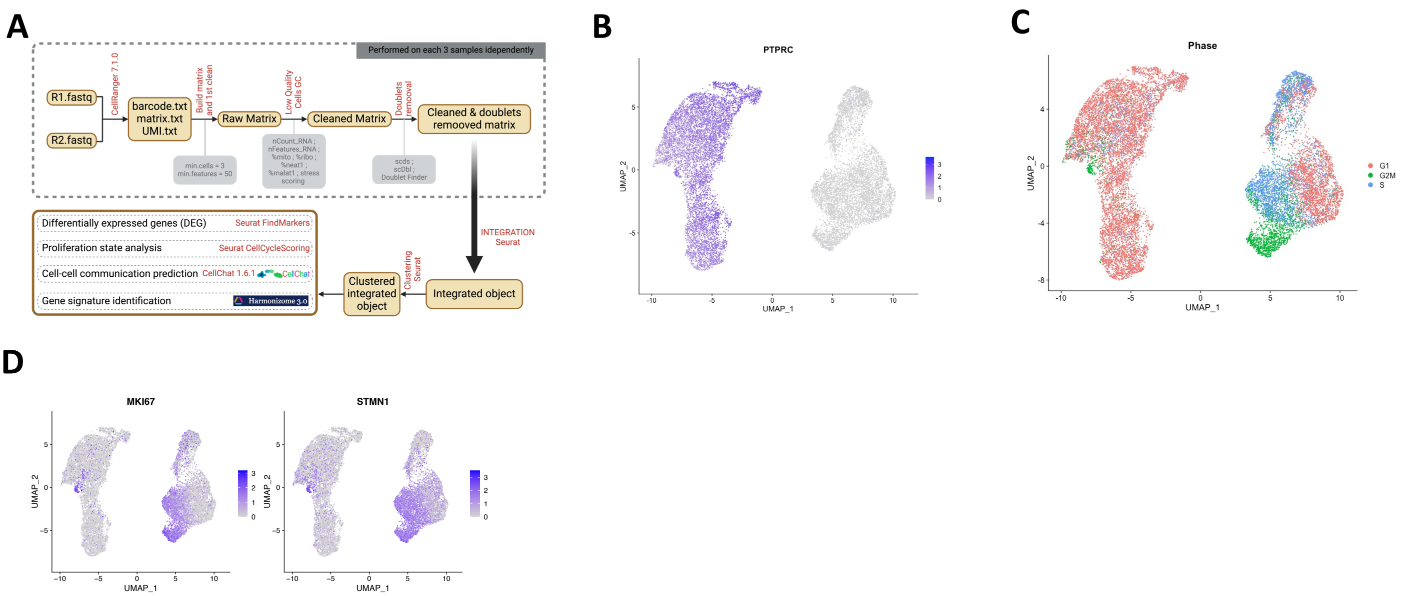

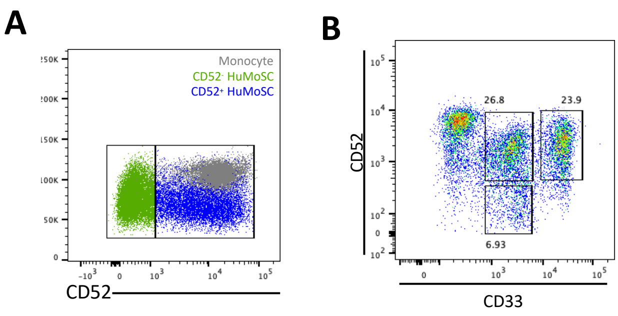

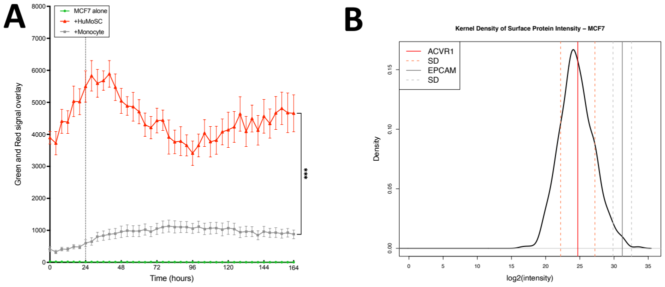

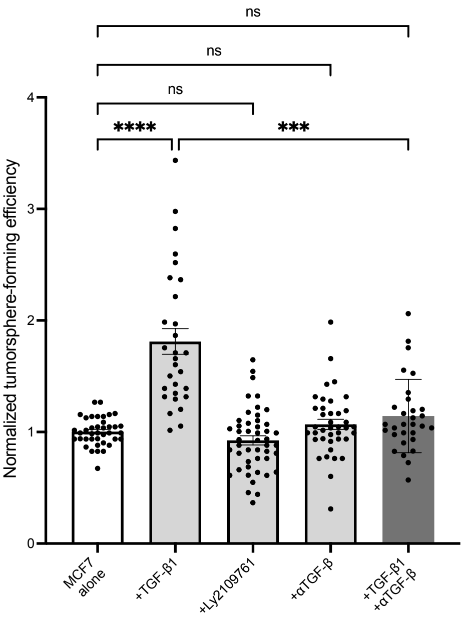
